# Vulnerability of ARID1A deficient cancer cells to pyrimidine synthesis blockade

**DOI:** 10.1101/2020.10.12.331975

**Authors:** Zhigui Li, Shijun Mi, Oloruntoba I. Osagie, Jing Ji, Chia-Ping H. Yang, Melissa Schwartz, Pei Hui, Gloria S. Huang

## Abstract

Here we report the discovery and preclinical validation of a novel precision medicine strategy for *ARID1A*-mutated cancer. Unbiased proteomics reveals for the first time that ARID1A protein (BAF250a) binds aspartate transcarbamoylase (ATCase), a key regulatory enzyme of the *de novo* pyrimidine synthesis pathway. Using isogenic paired ARID1A proficient/deficient cancer cell lines, we show that ARID1A protein deficiency (as occurs in *ARID1A* mutant cancers) leads to metabolic reprogramming and pyrimidine synthesis dependency. Pyrimidine synthesis blockade using the FDA-approved drug teriflunomide (a DHODH inhibitor) suppresses tumor growth and selectively induces DNA damage in ARID1A-deficient tumor models. Combining pyrimidine synthesis inhibition with DNA damage repair blockade, using teriflunomide and AZD6738 (an ATR inhibitor), achieves potent synergy and induces sustained tumor regression in *ARID1A*-mutant ovarian cancer patient-derived xenografts (PDX). These compelling preclinical data support the evaluation of this novel combination treatment in patients with *ARID1A*-mutated cancers.

**SIGNIFICANCE:** We identified that ARID1A-deficient cells are selectively vulnerable to pyrimidine synthesis blockade. Preclinical studies demonstrate the *in vivo* efficacy of a synergistic drug combination that concurrently inhibits the de novo pyrimidine synthesis pathway and DNA damage repair to induce regression in patient-derived xenograft models of ARID1A-mutated cancer.

## INTRODUCTION

*ARID1A* is among the most commonly mutated tumor suppressor genes in human cancer. The highest frequency of *ARID1A* mutations occur in gynecologic malignancies, including clear cell ovarian carcinoma (46-57%), endometrioid ovarian carcinoma (30%), uterine endometrial carcinoma (34%) and uterine carcinosarcoma (20-36%)^1–3^. *ARID1A* mutations are also observed in ~10-20% of diverse cancer types including gastric carcinoma, cholangiocarcinoma, bladder urothelial carcinoma, hepatocellular carcinoma, esophageal adenocarcinoma, cutaneous melanoma, and colorectal carcinoma. Mutations in *ARID1A* are strongly correlated with loss of protein expression^1,4^. The protein encoded by *ARID1A* is a core subunit of the BAF (mammalian SWI/SNF) chromatin remodeling complex, which modulates gene expression by binding to AT-rich DNA regions, mobilizing nucleosomes, and interacting with transcription factors, coactivators, and corepressors^5^. Among BAF complex subunits, *ARID1A* is the most frequently mutated in human cancer.

Patients with *ARID1A* mutated cancers have worse clinical outcomes compared to patients with *ARID1A* wildtype cancers. Overall survival is significantly shorter in patients with *ARID1A* mutated cancers compared to *ARID1A* wildtype cancers, when analyzing a pan-cancer cohort and when analyzing cancer-specific cohorts of patients with ovarian, hepatocellular, or pancreatic cancer^6^. Loss of ARID1A is also linked to shorter progression-free survival and chemoresistance^7^. The high frequency of *ARID1A* mutations in human cancer, and the unmet need for effective treatment for ARID1A deficient cancers, led us to undertake this study to uncover novel ARID1A functions associated with targetable therapeutic vulnerabilities.

In this study, we used an unbiased proteomics approach to identify novel protein-protein interactions of ARID1A. We show that ARID1A directly binds to ATCase, one of three key regulatory enzymes of the *de novo* pyrimidine synthesis pathway encoded by the *CAD* gene (Carbamoyl-Phosphate Synthetase 2, Aspartate Transcarbamylase, And Dihydroorotase). ARID1A protein deficiency (as occurs in *ARID1A*-mutated cancers) leads to metabolic reprogramming characterized by an increased rate of *de novo* pyrimidine synthesis and selective vulnerability to pyrimidine synthesis blockade, thus identifying an Achilles’ heel that can be therapeutically exploited. Furthermore, combination treatment with pyrimidine synthesis blockade and ataxia telangiectasia and rad3-related (ATR) kinase inhibition is potently synergistic and induces tumor regressions *in vivo*. Our results show a novel targetable function of ARID1A as a regulator of *de novo* pyrimidine synthesis and provide a therapeutic strategy to exploit the dependency of ARID1A deficient tumors on this metabolic pathway.

## RESULTS

### ARID1A interacts with the ATCase domain of CAD

To investigate unknown functions of ARID1A, we used mass spectrometry to analyze the immunoaffinity-purified ARID1A complex with the aim to identify novel ARID1A-interacting proteins. First, the endogenous ARID1A complex was immunoprecipitated from *ARID1A* wildtype endometrial cancer cells (KLE)^8^. The resulting Coomassie stained SDS-PAGE gel is shown in Fig. 1a. In addition to known BAF complex proteins, a ~250 kD protein band was observed and excised for analysis. Following trypsin proteolysis, the peptide sequences were determined by liquid chromatography-tandem mass spectrometry (LC-MS/MS) and identified to be Carbamoyl-phosphate synthetase 2, Aspartate transcarbamoylase, and Dihydroorotase (CAD). Fragmentation spectra of CAD peptides is shown in Supplementary Fig. S1a. We also performed LC-MS/MS analysis of the immunopurified ARID1A complex from *ARID1A* wildtype ovarian cancer cells (ES2)^9^ and obtained similar results identifying CAD peptides. Based on functional similarities analysis (Supplementary Fig. S1b), CAD differs from known members of the ARID1A interactome (i.e. BAF complex proteins) and thus is newly identified to be an ARID1A-interacting protein.

**Figure 1.**
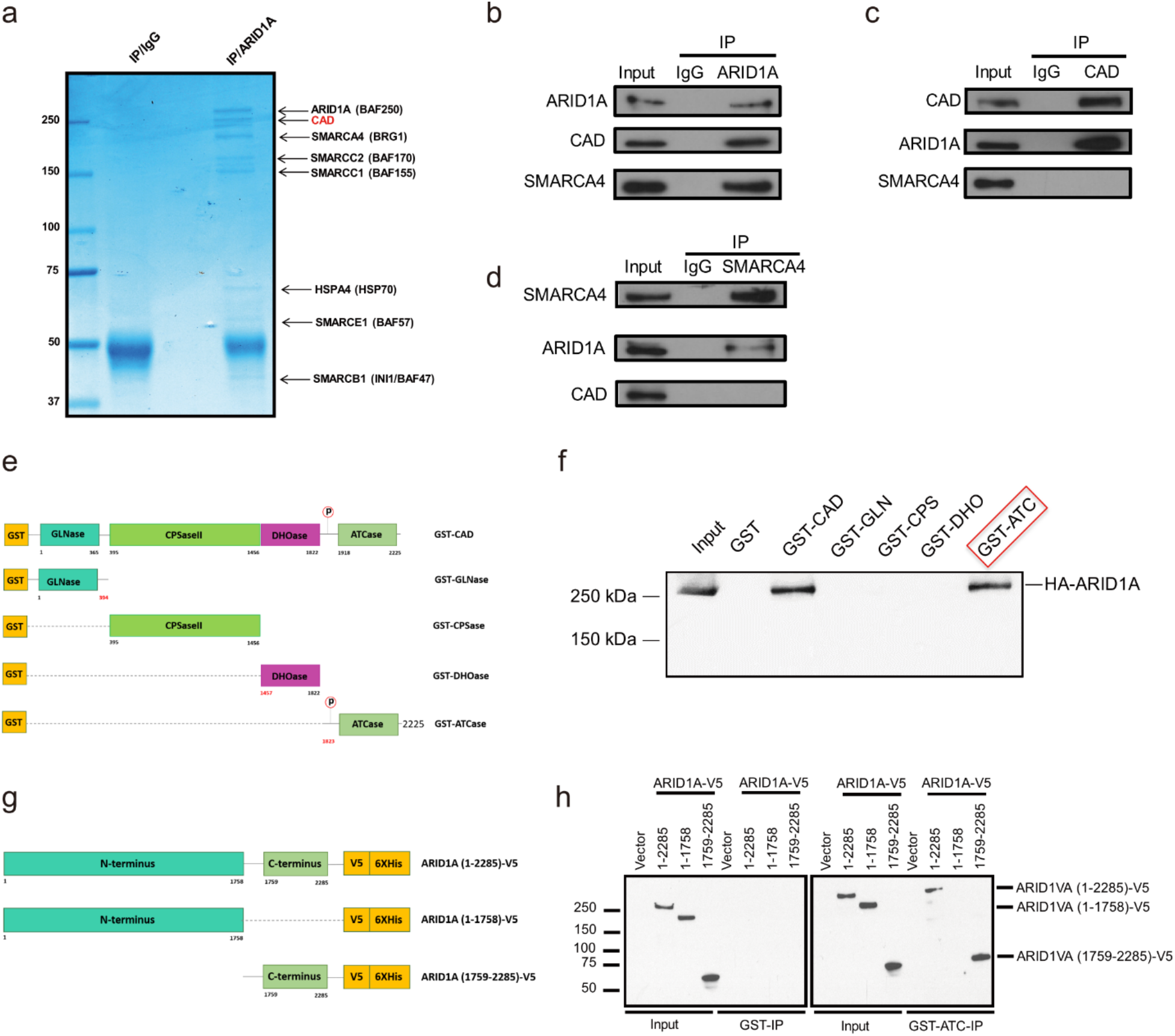
ARID1A interacts with CAD. **a**, Coomassie blue staining of ARID1A complex immunopurified from an *ARID1A*-wildtype cell line (KLE) is shown in the right lane, and immunopurified control IgG complex is shown in the left lane. Arrows designate protein band identification by LC-MS/MS, with black text for ARID1A and SWI/SNF family members and red text for the multifunctional enzyme CAD. **b-d**, Endogenous protein-protein interactions in an *ARID1A*-wildtype cell line (ES2) were assessed. Immunoprecipitation with the indicated antibody or control IgG antibody was performed, and immunoprecipitated proteins were detected by immunoblotting. At least two independent experiments were done, and representative immunoblots are shown. In (**b**), immunoprecipitation was done using an anti-ARID1A antibody. Both CAD and the core SWI/SNF subunit SMARCA4 co-immunoprecipitate with ARID1A. In (**c**), immunoprecipitation was done using an anti-CAD antibody. Endogenous ARID1A, but not SMARCA4, co-immunoprecipitates with CAD. In (**d**), immunoprecipitation was done using an anti-SMARCA4 antibody. Endogenous ARID1A, but not CAD, co-immunoprecipitates with SMARCA4. **e**, Recombinant, glutathione S-transferase (GST)-tagged proteins were made to express full-length CAD or one of four unique non-overlapping CAD fragments. Each of the CAD fragments contains a functional enzyme component, as shown. CAD fusion proteins were expressed in bacteria. **f**, Whole-cell lysates were prepared using HEK293T cells made to express HA-tagged full-length ARID1A. Shown are the results of a GST pulldown assay using recombinant GST-CAD fusion proteins (shown in A), followed by immunoblotting using an anti-HA antibody to detect HA-ARID1A. Recombinant full-length GST-CAD and the GST-ATCase domain demonstrated *in vitro* binding to HA-tagged ARID1A. **g**, Recombinant ARID1A-V5 fusion proteins were made for expression of full-length ARID1A (1-2285), N-terminal ARID1A (1-1758), or C-terminal ARID1A (1759-2285) in HEK293T cells. **h**, The GST-ATCase fusion protein (but not the control GST protein assessed on the left side of the panel) demonstrated *in vitro* binding to V5-tagged, full-length ARID1A (1-2285) and to C-terminal ARID1A (1759-2285), but not to N-terminal ARID1A (1-1758). These data indicate that the protein-protein interaction of ARID1A and CAD is localized to the C-terminal regions of both CAD (ATCase domain, 1823-2225) and ARID1A (1759-2285).

Immunoprecipitation using anti-ARID1A (Fig. 1b) and anti-CAD (Fig. 1c) antibodies followed by immunoblotting confirms the interaction between endogenous ARID1A and CAD in cell lines that express wild-type *ARID1A*. Co-immunoprecipitation results from *ARID1A*-wildtype ovarian cancer (ES2)^8^ cells are shown in Fig. 1. Similar results are observed in *ARID1A*-wildtype endometrial cancer cells (KLE), data not shown. In addition to ARID1A, a core BAF complex subunit is SMARCA4 (also known as BRG1)^10^. As expected, SMARCA4 co-immunoprecipitates with ARID1A (Fig. 1b). In contrast, SMARCA4 does not co-immunoprecipitate with CAD, nor does CAD co-immunoprecipitate with SMARCA4 (Fig. 1c, 1d). ARID1A’s interaction with CAD appears to be distinct from its known role as a BAF complex member.

The nature of the protein-protein interaction of CAD and ARID1A was further investigated. Recombinant GST-tagged full-length CAD and CAD protein fragments, corresponding to the protein domains shown in Fig.1e, were expressed in *Escherichia coli*. The *in vitro* interaction of full-length CAD with ARID1A was demonstrated by GST pulldown assay (Fig. 1f). By using recombinant GST-tagged CAD fragments as bait, the ARID1A-interacting domain of CAD is localized to the aspartate transcarbamylase (ATCase) domain (Fig. 1f). In the next set of experiments, GST-ATCase was used as the bait, and full-length ARID1A, N-terminus ARID1A, or C-terminus ARID1A was expressed in HEK293 (Fig. 1g). GST-ATCase pulls down full-length ARID1A, as expected, and C-terminus ARID1A but not N-terminus ARID1A (Fig. 1h). Thus, ATCase is the ARID1A-interacting domain of CAD, and the C-terminus of ARID1A binds to ATCase.

### ARID1A is a negative regulator of *de novo* pyrimidine biosynthesis

Next, we investigated the role of ARID1A as a potential regulator of CAD via its protein-protein interaction. *ARID1A* mutations in human cancers are typically associated with loss of ARID1A (BAF250A) protein expression due to truncating nonsense or frameshift mutations. To interrogate the functional consequences of ARID1A mutation and protein deficiency, we used previously validated short hairpin (sh) RNA vectors^11^ to knockdown *ARID1A*, and expanded stably transfected clones, called shARID1A(a) and shARID1A(b), for further analysis. Knockdown of *ARID1A* was confirmed by immunoblotting (Fig. 2a and 2b). The resulting phenotype was evaluated relative to isogenic cells transfected with a non-targeting scrambled control shRNA (shCon) and untransfected cells.

**Figure 2.**
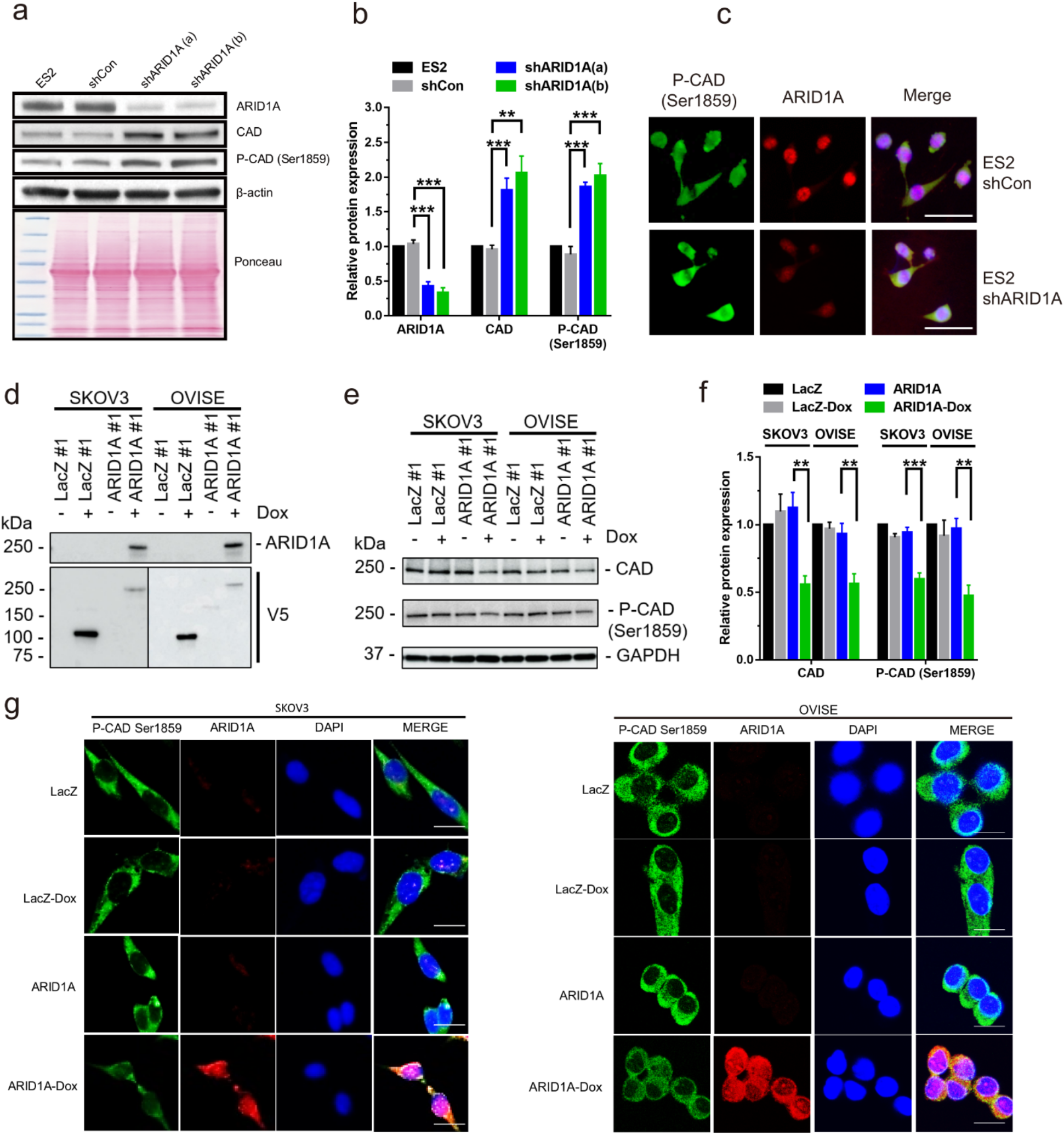
ARID1A is a negative regulator of CAD. **a**, Immunoblotting using an anti-ARID1A antibody shows knockdown of *ARID1A* protein expression following stable transfection with short hairpin RNAs, shARID1A(a) and shARID1A(b), compared with control transfection with a non-targeting short hairpin RNA, shCon. Protein expression levels of total CAD and phosphorylated CAD increased in the *ARID1A*-knockdown cells compared to control cells, as shown by immunoblotting with anti-CAD and anti-phosphorylated-CAD antibodies. Equal protein loading is shown by immunoblotting with an anti-β-actin antibody and Ponceau staining of the nitrocellulose membrane. At least five independent experiments were done, and representative results are shown. **b**, Quantitation of immunoblotting images in **a** by Image J. Data represent mean ± s.e.m, *n* = 3 independent experiments. One-way ANOVA with Tukey’s post-test; \*\**P* < 0.01, \*\*\**P* < 0.001. **c**, Representative immunofluorescence staining of ES2 cells for phosphorylated CAD Ser1859 (green) and ARID1A (red) is shown. P-CAD Ser1859 protein increased in the *ARID1A*-knockdown cells compared to control cells. Nuclei are indicated by DAPI staining. Scale bar, 40 μm. **d**, Immunoblotting shows restoration of ARID1A protein expression following stable transfection with Lenti-puro-ARID1A-V5 (ARID1A) compared with control transfection with non-targeting Lenti-puro-LacZ-V5 (LacZ). Protein expression levels of ARID1A and V5 tag in the ARID1A-restoration cells compared to control cells were shown by immunoblotting with anti-ARID1A and anti-V5 antibodies. At least three independent experiments were done, and representative results are shown. **e**, To evaluate the effect of ARID1A on CAD protein, immunoblotting using anti-CAD and anti-CAD Ser1859 antibodies was performed. CAD and phosphorylated CAD protein expression decreased following stable transfection with Lenti-puro-ARID1A (ARID1A) compared with control transfection with non-targeting Lenti-puro-LacZ (LacZ). Equal protein loading is shown by immunoblotting with an anti-GAPDH antibody. At least three independent experiments were done, and representative results are shown. **f**, Quantitation of immunoblotting images in **d** by Image J. Data represent mean ± s.e.m, *n* = 3 independent experiments. One-way ANOVA with Tukey’s post-test; \*\**P* < 0.01, \*\*\**P* < 0.001. **g**, Representative immunofluorescence staining of the SKOV3 (left) and OVISE (right) cell lines for phosphorylated CAD Ser1859 (green) and ARID1A (red) is shown. P-CAD Ser1859 protein decreased in the ARID1A-restoration cells following induction by doxycycline (Dox) for 2 days compared to control cells. Nuclei are indicated by DAPI staining. Scale bar, 20 μm.

As shown in Fig. 2a and 2b, *ARID1A* knockdown cells demonstrate higher total CAD levels relative to control cells. CAD activity is controlled via phosphorylation at key regulatory sites including serine 1859. This CAD site is phosphorylated by ribosomal protein S6 kinase B1 (RPS6KB1, also known as S6K)^12,13^. Shown in Fig. 2a and 2b, serine 1859 phosphorylated CAD protein (P-CAD Ser1859) levels increase following *ARID1A* knockdown. In *ARID1A* knockdown cells, RPS6KB1 activity is similar as in control cells; evaluation of the RPS6KB1 canonical substrate, ribosomal protein S6 by immunoblotting is shown in Supplementary Fig. S2. Thus, total and phosphorylated CAD levels following *ARID1A* knockdown is not associated with altered RPS6KB1 activity and is inversely correlated with the ARID1A protein level.

The inverse correlation of ARID1A levels with CAD levels was confirmed by immunofluorescence analysis in several independent isogenic paired cell lines, and representative data is shown in Fig. 2c. Following *ARID1A* knockdown, the protein expression of phosphorylated CAD level is increase compared with control cells.

Complementing these experiments, the effect of ARID1A restoration on CAD was determined using *ARID1A*-mutant ovarian cancer cell lines, SKOV3 and OVISE (Supplementary Table S10). Both of these cell lines contain *ARID1A* mutations, associated with loss of ARID1A protein expression. To restore *ARID1A* expression, SKOV3 cells and OVISE cells were stably transfected with a tetracycline inducible system to express full-length wildtype *ARID1A* and compared with empty vector control. Restored ARID1A protein expression was confirmed by immunoblotting (Fig. 2d). The total and phosphorylated CAD protein levels decrease in *ARID1A* restoration cells (Fig. 2e and 2f). The inverse correlation of ARID1A and phosphorylated CAD levels in the *ARID1A* restoration cell lines is also evident by immunofluorescence (Fig. 2g).

Similarly, ARID1A restoration was done in HEC-1-A, an *ARID1A*-mutant endometrial cancer cell line. This cell line has two heterozygous truncating mutations at p.Q1835* and p.Q2115* associated with loss of ARID1A protein expression^14^. Following transfection with full-length ARID1A, compared with control empty vector, total and phosphorylated CAD levels decrease (Supplementary Fig. S3).

Collectively, these findings indicate that wildtype ARID1A functions as a negative regulator of CAD, wherein ARID1A deficiency results in an increase in total and phosphorylated CAD.

### ARID1A deficiency promotes *de novo* pyrimidine biosynthesis

The ATCase enzymatic activity of CAD catalyzes the reaction of carbamoyl phosphate and aspartate to N-carbamoyl aspartate, and is the first committed step in *de novo* pyrimidine biosynthesis^15^ (Fig. 3a). We next investigated whether ARID1A deficiency affects *de novo* pyrimidine synthesis. To quantify the rate of *de novo* pyrimidine synthesis, incorporation of ^14^C-radiolabelled aspartate into RNA and DNA was measured. RNA and DNA synthesized via the pyrimidine salvage pathway do not incorporate the ^14^C-radiolabelled aspartate, imparting specificity for measuring *de novo* pyrimidine synthesis flux. We found that *ARID1A* knockdown results in increased ^14^C incorporation into RNA, indicating increased flux through the *de novo* pyrimidine synthesis pathway (Fig. 3b). Increased ^14^C incorporation is similarly observed in DNA (data not shown). This demonstrates the inverse relationship of ARID1A levels and the rate of *de novo* pyrimidine synthesis.

**Figure 3.**
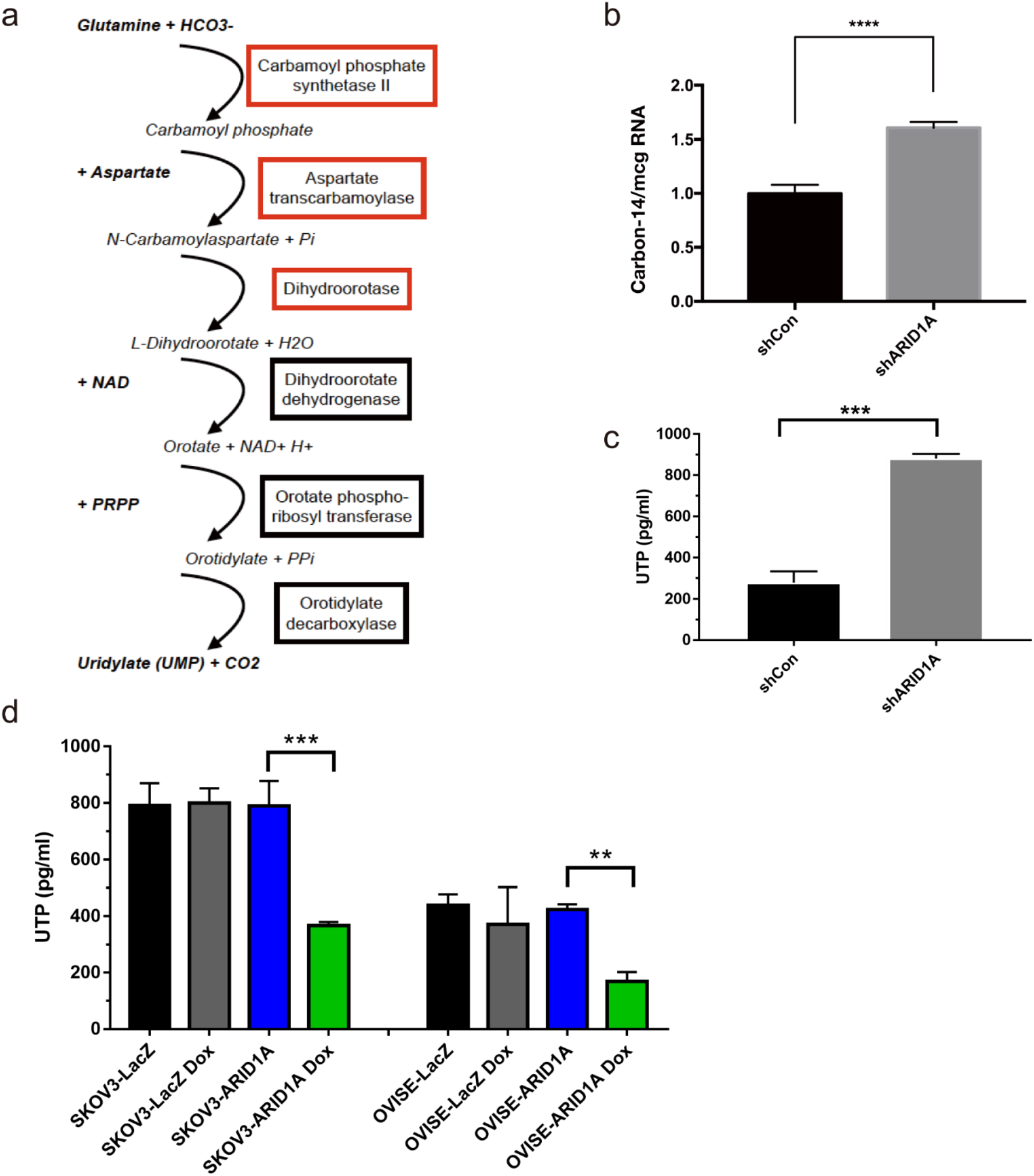
ARID1A deficiency promotes *de novo* pyrimidine synthesis. **a**, The diagram shows the steps in the *de novo* pyrimidine synthesis pathway. Aspartate is incorporated by aspartate transcarbamoylase in the second step of *de novo* pyrimidine synthesis. **b**, To evaluate the effect of *ARID1A* knockdown on *de novo* pyrimidine synthesis, carbon-14-labeled aspartate was added to the growth media. After a 6-hour incubation, RNA was isolated from the cells, and carbon-14 incorporation into the RNA was quantified as the counts per minute (cpm) measured by a scintillation counter and was normalized to the amount of RNA (micrograms). The bar graph shows increased carbon-14 incorporation into RNA in *ARID1A*-knockdown cells relative to control cells. The mean ± SD calculated from two independent experiments, each performed in two to four biological replicates, is shown. \*\*\*\**P* < 0.0001, two-tailed *t*-test. **c**, The cellular UTP level was evaluated in ARID1A-knockdown cells. UTP increased in *ARID1A*-knockdown ES2 cells relative to control cells transfected with a non-targeting short hairpin RNA, shCon. The UTP level in ES2 parental and another *ARID1A*-Knockdown ES2 cells show similar result (Fig. S3a). Teriflunomide (Teri) treatment decreased UTP in the ES2 panel (Fig. S3b). Differences in UTP were evaluated using one-way ANOVA with Tukey’s post-test; \*\*\**P* < 0.001. **d**, UTP decreased in ARID1A-restoration cell lines (SKOV3, left, and OVISE, right) following induction by doxycycline (Dox) compared with control cells transfected with non-targeting Lenti-puro-LacZ (LacZ). Differences in UTP were evaluated using one-way ANOVA with Tukey’s post-test; \*\**P* < 0.01, \*\*\**P* < 0.001.

An elevated rate of *de novo* pyrimidine biosynthesis may result in higher steady state levels of its product Uridine-5’-triphosphate (UTP). Thus, we examined UTP levels in *ARID1A* knockdown and restoration cell line panels. The UTP level is increased in *ARID1A* knockdown cells compared with control *ARID1A* wildtype cells (Fig. 3c and Supplementary Fig. S4a). The UTP level is reduced in *ARID1A* restoration cells compared with control *ARID1A*-mutant cells, SKOV3 and OVISE (Fig. 3d). Together, these data indicate that the interaction of ARID1A and ATCase regulates *de novo* pyrimidine biosynthesis flux and consequently, the pyrimidine nucleotide pool.

### *ARID1A*-deficient cells and tumors display sensitization to *de novo* pyrimidine synthesis blockade therapy

We evaluated the effect of FDA-approved inhibitors of dihydroorotate dehydrogenase (DHODH), the enzyme immediately downstream of CAD that catalyzes the conversion of dihydroorotate to orotate. As shown in Fig. 4a, *ARID1A* knockdown cells are significantly more sensitive to the DHODH inhibitor teriflunomide. Similar results are observed with the DHODH inhibitor leflunomide (data not shown). *ARID1A*-mutant ovarian cancer cell lines SKOV3 and OVISE are sensitive to teriflunomide, while *ARID1A* restoration decreases sensitivity (Fig. 4b and 4c).

**Figure 4.**
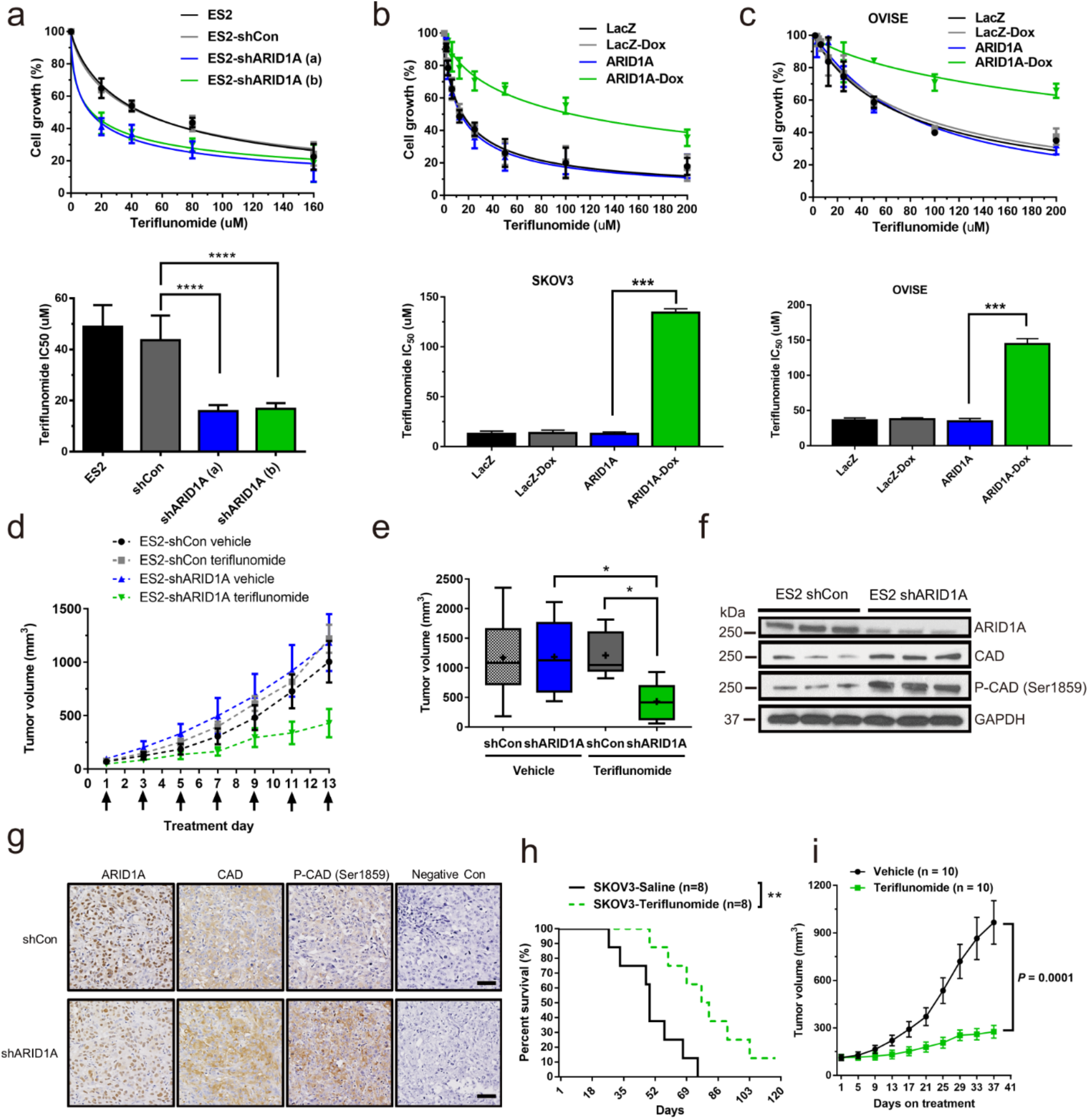
*ARID1A*-deficient cells and tumors display sensitization to *de novo* pyrimidine synthesis blockade therapy. **a**, In the upper panel, the effect of teriflunomide in ES2 cells is quantified by showing the relative cell number following drug treatment at various concentrations for 72 h. *ARID1A*-knockdown cells, depicted by the blue line for shARID1A (a) and the green line for shARID1A (b), were more sensitive to teriflunomide, resulting in a decreased cell number following drug treatment, compared to untransfected cells (black line) or shCon cells (gray line). In the lower panel, the data shown in the upper panel was summarized by graphing the teriflunomide concentration that results in a 50% growth inhibitory effect (IC_50_). The bars depict the mean IC_50_ ± SD for five independent experiments. Differences in IC_50_ were evaluated using one-way ANOVA with Tukey’s post-test; \*\*\*\**P* < 0.001. **b**, In the upper panel, the effect of teriflunomide is quantified in a single-clone SKOV3 cell line that was stably transfected with Lenti-puro-ARID1A-V5 (ARID1A) compared with control cells transfected with non-targeting Lenti-puro-LacZ-V5 (LacZ). ARID1A-restoration cells induced by doxycycline (Dox) (green line) were more resistant to teriflunomide, resulting in an increased cell number following drug treatment, compared to uninduced cells (blue line). In the lower panel, the data shown in the upper panel was summarized by graphing the IC_50_. The bars depict the mean IC_50_ ± SD for three independent experiments. Differences in IC50 were evaluated using one-way ANOVA with Tukey’s post-test; \*\*\**P* < 0.001. **c**, In the upper panel, the effect of teriflunomide is quantified in a single-clone OVISE cell line that was stably transfected with Lenti-puro-ARID1A-V5 (ARID1A) compared with control cells transfected with non-targeting Lenti-puro-LacZ-V5 (LacZ). ARID1A-restoration cells induced by Dox (green line) were more resistant to teriflunomide, resulting in an increased cell number following drug treatment, compared to uninduced cells (blue line). In the lower panel, the data shown in the upper panel was summarized by graphing the IC_50_. The bars depict the mean IC_50_ ± SD for three independent experiments. Differences in IC_50_ were evaluated using one-way ANOVA with Tukey’s post-test; \*\*\**P* < 0.001. **d**, The effect of teriflunomide was evaluated *in vivo* by treating xenograft-bearing mice every other day for 13 days (black arrows). The effect of teriflunomide on tumor xenograft growth is shown by depiction of the mean tumor volume ± SD (N=6 to 8 animals/group) for each of the following groups: shCon treated with vehicle (black line), shARID1A treated with vehicle (blue line), shCon treated with teriflunomide (gray line), and shARID1A treated with teriflunomide (green line). Teriflunomide effectively inhibited tumor growth of shARID1A xenografts (green line), but not shCon xenografts (gray line). **e**, The data shown in (d) are summarized by graphing the terminal tumor volumes. The bars depict the median (middle line) and mean (+) tumor volumes for each group. Differences between groups were evaluated using one-way ANOVA with Tukey’s post-test; \**P* < 0.05. **f**, ARID1A, CAD, and phospho-CAD Ser1859 protein expression was evaluated in representative tumor xenograft cell lysates by immunoblotting. *ARID1A*-knockdown xenografts (shARID1A) showed decreased ARID1A protein levels and increased CAD and phospho-CAD protein levels. Equal protein loading was confirmed by Ponceau staining of the nitrocellulose membrane (not shown) and immunoblotting with an anti-GAPDH antibody. **g**, Representative immunohistology staining of xenograft tumor tissue sections for ARID1A, CAD, and phosphorylated CAD Ser1859 is shown. The negative-control samples underwent the same immunohistology staining procedure but without primary antibody incubation. Similar upregulation of CAD and P-CAD Ser1859 was observed in the *ARID1A*-knockdown xenograft tumor samples. Scale bar, 50 μm. **h**, The effect of teriflunomide was evaluated in an *ARID1A*-deficient SKOV3 tumor xenograft model. The xenograft model was generated by subcutaneously injecting SKOV3 cells in Matrigel (1:1) into athymic nude mice. Teriflunomide (4 mg/kg) or vehicle was intraperitoneally injected every other day. Tumor size was recorded on the same day. A survival curve is shown; the terminal tumor volume is 1000 mm^3^. Teriflunomide improved the survival of the tumor-bearing mice. The *P* value was calculated via two-tailed *t*-test. \*\**P* < 0.01. The effect of teriflunomide on an ES2-shCon tumor xenograft model is shown in Figure S6. **i**, PDXs (CTG-2213; ARID1A truncating mutation at Gln211*) were randomized into vehicle control and Teriflunomide (4 mg/kg every other day). Tumor volume was measured every four days. There was a significant decrease in treatment group relative to control (*P* = 0.0001). The effect of teriflunomide on animal weight and individual tumor growth is shown in Supplementary Fig. S7.

Vulnerability of ARID1A deficient cells to *de novo* pyrimidine synthesis blockade was confirmed using isogenic *ARID1A* knockout and *ARID1A* wildtype HCT116 colorectal carcinoma cells. Homozygous deletion of *ARID1A* results from knock-in of premature stop codons (Q456*/Q456*). Compared to wildtype *ARID1A* cells, *ARID1A* knockout cells are hypersensitive to DHODH inhibitors (Supplementary Fig. S5).

*In vivo* validation is an important step in translation of scientific findings to clinical application. Therefore, we evaluated the therapeutic efficacy of DHODH inhibition using clear cell ovarian cancer xenografts (Fig. 4d). Xenograft-bearing mice were randomized to treatment with the DHODH inhibitor teriflunomide, or vehicle alone. Teriflunomide was administered intraperitoneally at a well-tolerated dosing regimen of 4 mg/kg every other day, corresponding to a human equivalent dose of 0.32 mg/kg every other day. This dose level is ~30% lower than the FDA-approved dose level of 14 mg daily used for treating multiple sclerosis^16^. As shown in Fig. 4d and 4e, teriflunomide selectively suppresses tumor growth in *ARID1A*-deficient xenografts, compared to ARID1A-wildtype xenografts (Supplementary Fig. S6). We also evaluated the effect of teriflunomide in *ARID1A*-mutant SKOV3 tumor xenograft models (Fig. 4h). Teriflunomide significantly improves animal survival compared to the vehicle treatment group.

Patient-derived xenograft (PDX) models are particularly valuable for preclinical validation. We used an *ARID1A*-mutant clear cell ovarian cancer PDX model (CTG-2213; ARID1A truncating mutation at Gln211*) developed from direct implantation of fresh viable human tumor tissue propagated in suitable mouse hosts. PDX models accurately recapitulate tumor heterogeneity and predict clinical response to therapy. Shown in Fig. 4i, tumor volume is significantly reduced in the teriflunomide treatment group relative to the vehicle control group (*P* < 0.001).There was no weight loss or toxicity observed in mice in either the treatment or vehicle control groups (Supplementary Fig. S7). These data demonstrate the selective *in vivo* efficacy of DHODH inhibition in multiple *ARID1A*-deficient cancer models.

### DHODH inhibitor therapy induces DNA damage repair

To investigate ARID1A-dependent differences in the cellular response to DHODH inhibition, we evaluated DNA damage following drug treatment of isogenic ARID1A-proficient and deficient cells. As shown by gamma-H2AX immunofluorescence and immunoblotting (Fig. 5i-k), teriflunomide selectively induces DNA damage in ARID1A-deficient cells relative to ARID1A-proficient ES2 cells. Teriflunomide treatment activates the CHK1 DNA repair pathway, as shown by a robust increase in Ser-345 phosphorylation of CHK1 in ES2-shARID1A cells compared with the ES2-shCon cells (Fig. 5i-j). These results indicate that the cellular response of DHODH inhibition depends on the ARID1A status. In *ARID1A*-deficient cells, induction of DNA damage by teriflunomide triggers DNA damage repair signaling pathways that involve CHK1 kinase activation.

**Figure 5.**
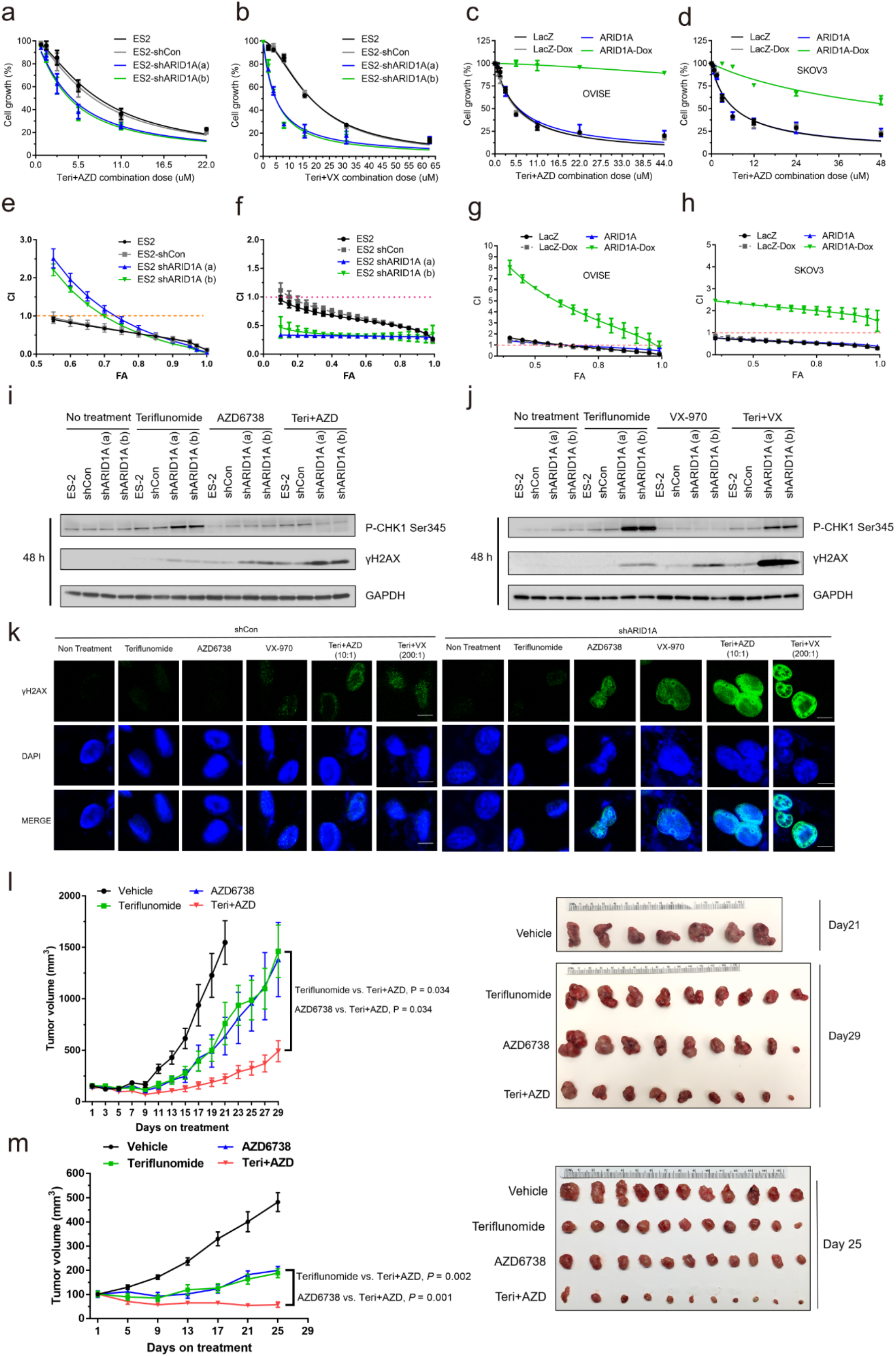
Combination treatment with DHODHi and ATRi shows a synergistic effect in *ARID1A*-deficient cells. **a**, The effect of combination treatment of ES2 cells with teriflunomide and AZD6738 for 72 h is quantified. *ARID1A*-knockdown cells, depicted by the blue line for shARID1A (a) and the green line for shARID1A (b), were even more sensitive to combination treatment compared to untransfected cells (black line) or shCon cells (gray line). Representative images of cell morphology at 72 h are shown in Supplementary Figure S8. **b**, The effect of combination treatment of ES2 cells with teriflunomide and VX-970 for 72 h is quantified. *ARID1A*-knockdown cells, as depicted by the blue line for shARID1A (a) and the green line for shARID1A (b), are even more sensitive to combination treatment compared to untransfected cells (black line) or shCon cells (gray line). **c**, The effect of combination treatment of ARID1A-restoration OVISE cells with teriflunomide and AZD6738 for 72 h is quantified. ARID1A-induced cells (ARID1A-Dox), depicted by the green line, were even more resistant to combination treatment compared to non-ARID1A-induced cells. **d**, The effect of combination treatment of ARID1A-restoration SKOV3 cells with teriflunomide and AZD6738 for 72 h is quantified. ARID1A-induced cells (ARID1A-Dox), depicted by the green line, were even more resistant to combination treatment compared to non-ARID1A-induced cells. **e-h**, The data shown in (**a-c**) are summarized by graphing FA-CI plots, with × = fraction affected (FA) vs. y = combination index (CI) (the Chou-Talalay plot). The FA-CI curves showed the detail synergic effect at each drug concentration. CI < 1, = 1, and > 1 indicate synergism, an additive effect, and antagonism, respectively. Detailed analyses of the combination treatments in (**a**), (**b**), (**c**) and (**d**) are shown in Supplementary Tables S1, S5, S6 and S7, respectively. **i**, γ-H2AX protein expression was evaluated in ES2 cell lysates by immunoblotting at both 48 h and 72 h. Teriflunomide (15 μM) and AZD6738 (1.5 μM) were used in single-drug and combination treatment groups. The target effect of AZD6738 on P-CHK1 Ser345 was decreased. *ARID1A*-knockdown cells (shARID1A) in the single-drug treatment groups showed increased γ-H2AX protein levels, but the highest levels were in the Teri+AZD combination treatment group. Equal protein loading was confirmed by immunoblotting with an anti-GAPDH antibody. **j**, The Teriflunomide+VX-970 combination treatment effect was evaluated by detecting γ-H2AX protein expression in ES2 cell lysates by immunoblotting at both 24 h and 48 h. The target effect of VX-970 on P-CHK1 Ser345 was decreased. Teriflunomide (15 μM) and VX-970 (0.075 μM) were used in single-drug and combination treatment groups. *ARID1A*-knockdown cells (shARID1A) showed increased γ-H2AX protein levels in the single-drug treatment groups, but the highest levels were in the combination treatment group. Equal protein loading was confirmed by immunoblotting with an anti-GAPDH antibody. **k**, Representative immunofluorescence staining of ES2 cells for γ-H2AX (green) and DAPI (blue) at 24 h. Teriflunomide (15 μM), AZD6738 (1.5 μM), and VX-970 (0.075 μM) were used in single-drug and combination treatment groups. *ARID1A*-knockdown cells (shARID1A) showed increased γ-H2AX protein levels following single-drug treatment groups, but the highest levels were in the combination treatment group. Scale bar, 10 μm. **l**, The effect of combination treatment with teriflunomide and AZD6738 was evaluated *in vivo* by treating ES2-shARID1A xenograft-bearing mice. The effect on tumor xenograft growth is shown by depiction of the mean tumor volume ± s.e.m. (N=7 to 9 animals/group) for each of the following groups: vehicle (black line, N=7), teriflunomide (green line, N=9), AZD6738 (blue line, N=9), and teriflunomide and AZD6738 (red line, N=9). Teriflunomide (4 mg/kg) or vehicle was intraperitoneally injected every other day. AZD6738 (25 mg/kg) or vehicle was given by oral gavage every day. Combination treatment more effectively inhibited tumor growth of shARID1A xenografts (red line) compared to single-drug-treated xenografts (green and blue lines). Images of all ES2-shARID1A xenografts treated with vehicle control, teriflunomide, AZD6738, or teriflunomide plus AZD6738 is shown on the right side. The images are from the endpoint of scheduled treatment. The vehicle group reached maximum tumor on day 21. Tumor size on day 29 showed that Combination treatment more effectively inhibited tumor growth of shARID1A xenografts compared to single-drug-treated xenografts. Analysis of the correlation of tumor weight with tumor volume in each group is shown in Supplementary Figure S11. **m**, The effect of combination treatment with teriflunomide and AZD6738 was evaluated in PDXs (CTG-2213; ARID1A truncating mutation at Gln211*). The effect on tumor xenograft growth is shown by depiction of the mean tumor volume ± s.e.m. (N=11 animals/group) for each of the following groups: vehicle (black line), teriflunomide (green line), AZD6738 (blue line), and teriflunomide plus AZD6738 (red line). Combination treatment more effectively inhibited tumor growth (red line) compared to single-drug-treated xenografts (green and blue lines). The images of all PDXs treated with vehicle control, teriflunomide, AZD6738, or teriflunomide plus AZD6738 on the right side. The images are from the endpoint of scheduled treatment on day 25. The combination treatment more effectively inhibited tumor growth of shARID1A xenografts compared to single-drug-treated xenografts. Analysis of the correlation of tumor weight with tumor volume in each group is shown in Supplementary Figure S11.

### ATR inhibition synergistically potentiates therapeutic effect of DHODH blockade

Since CHK1 kinase activation requires ATR activity, we hypothesized that ATR inhibition would prevent activation of protective DNA repair signaling and thereby enhance the efficacy of DHODH blockade.

*ARID1A*-deficient cells may rely on ATR-mediated DNA damage repair due to reduced activity of alternate DNA repair pathways^17^. We confirmed enhanced sensitivity to ATR inhibition in *ARID1A*-knockout HCT116 colorectal carcinoma cells compared to ARID1A-wildtype HCT116 cells (Supplementary Fig. S9). We also evaluated the ATR inhibitor response in ARID1A-knockout ES2 ovarian carcinoma cells compared to ARID1A-wildtype ES2. We report in a separate manuscript the *in vitro* and *in vivo* evaluation of ATR inhibitors (e.g. AZD-6738, VX-970) in multiple ovarian and endometrial models. Our results in isogenic models demonstrate that ARID1A deficiency confers sensitization to ATR inhibitors.

Next we evaluated the drug combination of teriflunomide with ATR inhibitors (AZD-6738, VX-970). Drug combination analysis by the method of Chou and Talalay^18^ demonstrates synergy of concurrent teriflunomide and ATR inhibition in multiple independent cancer cell lines (Fig. 5e-f; Supplementary Table S2-4). As shown using knockdown and CRISPR knockout experiments, *ARID1A*-deficient cells are significantly more sensitive to combination therapy compared to isogenic *ARID1A-*proficient cells (Fig. 5a-b; Supplementary Table S1, S5, S8; Supplementary Fig S10). *ARID1A* restoration in *ARID1A* mutant cells results in drug antagonism and diminishes the response to combination treatment (Fig. 5c-d, 5g-h; and Supplementary Table S6 and S7). As predicted, combination treatment results in potent induction of DNA damage in *ARID1A-*deficient cells compared to proficient cells (Fig. 5i-5k).

*In vivo* evaluation of Teriflunomide combined with ATR inhibitor AZD-6738 shows that the combination treatment is highly efficacious (Fig. 5l-m and Supplementary Fig. S11). Evaluated in two independent experiments using an aggressive ovarian cancer xenograft model (ES2-shARID1A) and an ARID1A-mutant clear cell ovarian cancer PDX model, the combination treatment is significantly more effective than single drug treatments (Fig. 5l-m). The animals maintain normal activity and weight throughout drug treatments which appear to be well tolerated. Sustained tumor regression is observed following combination treatment in the *ARID1A*-mutant clear cell ovarian cancer PDXs (Fig. 5m). Together, these data provide compelling preclinical data to support the efficacy of this novel combination treatment for *ARID1A*-mutant cancer.

## DISCUSSION

Our results reveal a novel therapeutically targetable function of ARID1A as a regulator of *de novo* pyrimidine synthesis. We show for the first time that ARID1A deficiency results in vulnerability to pyrimidine synthesis blockade.

We found that pyrimidine synthesis blockade using currently available FDA-approved drugs such as teriflunomide selectively suppresses cellular proliferation and induces DNA damage in *ARID1A*-deficient cells. *In vitro and in vivo* experiments demonstrate that ARID1A deficiency predicts sensitivity to teriflunomide. Based on these data, monotherapy with DHODH inhibitors may be useful for targeted treatment of cancers with *ARID1A* mutations.

The antitumor efficacy of DHODH inhibition is enhanced by concurrently exploiting the dependency of *ARIDA*-mutated cancers on ATR-mediated DNA repair. We show that ATR inhibitors synergize with teriflunomide to potentiate DNA damage and suppress cellular proliferation. This novel drug combination induced sustained tumor regression in a highly aggressive tumor model, *ARID1A*-mutated ovarian clear cell carcinoma PDXs. Combining pyrimidine synthesis blockade with DNA damage repair inhibitors is an attractive strategy for clinical evaluation in biologically aggressive *ARID1A*-mutated cancers.

We carried out protein interaction studies that show ARID1A directly binds to ATCase (one of three enzymes encoded by the *CAD* gene). ATCase, the primary regulatory enzyme of *de novo* pyrimidine biosynthesis, is allosterically regulated by ATP availability thereby functioning as a cellular energy sensor input to this pathway. In addition to its enzymatic activity being positively regulated by a purine (ATP), ATCase is negatively regulated by a pyrimidine (CTP), enabling its critical function of maintaining nucleotide pool balance between purines and pyrimidines^19,20^. Nucleotide pool balance is a key determinant of DNA replication fidelity. Thus, regulation of ATCase by ARID1A could play a role in maintaining nucleotide pool balance and DNA replication fidelity, and may contribute to ARID1A’s tumor suppressor function.

In summary, we discovered that ARID1A regulates the *de novo* pyrimidine synthesis pathway through an unexpected interaction with the energy-sensing enzyme ATCase. Metabolic reprogramming that results from ARID1A deficiency confers hypersensitivity to pyrimidine synthesis blockade, leading to therapeutic opportunities to repurpose FDA-approved inhibitors such as teriflunomide. Based on compelling data from *in vitro* and *in vivo* studies, we propose clinical trials of pyrimidine synthesis inhibitors alone and in combination with ATR inhibitors for precision therapy of *ARID1A*-mutated cancers.

## METHODS

### Reagents

#### Plasmids

HA-tagged full-length *ARID1A* was amplified by PCR from pCNA6-V5/His-ARID1A (provided by I.-M. Shih^11^) and subcloned into the pCIN4 expression vector^21^. To construct the expression plasmids for GST-CAD and GST-CAD fragments, cDNA sequences of full-length *CAD* and its fragments were amplified by PCR from pcDNA3.1-HisFlag-CAD (Addgene) and subcloned into the pGEX 4T-2 vector (GE Healthcare Life Sciences) for expression in BL21 bacteria. V5-tagged full-length BAF250a; V5-tagged BAF250a fragment, amino acids 1-1758; and V5-tagged BAF250a fragment, amino acids 1759-2285, were created by G. R. Crabtree^10^ and obtained from Addgene. Short hairpin RNA (shRNA) lentiviral plasmids were kindly provided by I.-M. Shih^8^. The shRNA sequences for *ARID1A* are as follows: sh1(TRCN0000059090), target sequence CCTCTCTTATACACAGCAGAT, and sh2(TRCN0000059091), target sequence CCGTTGATGAACTCATTGGTT. The vector backbone is pLKO.1. V5/His-tagged pLenti-puro-LacZ and pLenti-puro-ARID1A were obtained from Addgene.

#### Antibodies and Drugs

Antibodies for ARID1A, CAD, CAD (IHC), and β-actin were from Bethyl Laboratories. Antibodies for BRG1 (SMARCA4), Phospho-CAD (Serine 1859), Phospho-CHK1 (Serine 345), phosphor-Histone H2AX (γH2AX), HA, V5, RPS6, Phospho-RPS6, and GAPDH were from Cell Signaling Technology. Anti-IgG and HRP-labeled anti-rabbit secondary antibodies were from Invitrogen. Leflunomide was from Enzo Life Sciences. Teriflunomide and PF-4708671 were from Tocris. AZD6738 was from ChemScene. VX970 (VE-822) was from Selleck Chemicals.

#### Cell Lines

Adherent cell lines were cultured in RPMI 1640 (Gibco) supplemented with 10% fetal bovine serum (Gibco) and 1% penicillin-streptomycin (Gibco) at 37°C, 5% CO_2_. Low-passage-number cells were used, and all cell lines tested negative for mycoplasma using the MycoAlert Mycoplasma Detection Kit (Lonza). The following ovarian and endometrial cancer cell lines were used: ES2^9^, KLE^8^, SKOV3, OVISE (JCRB Cell Bank), and HEC-1-A^8^ cells and their stably transfected subclones, as described below. Cell lines were authenticated using the GenePrint10 kit (Promega) and matching to their original profiles (ATCC). ARID1A protein expression status was confirmed by immunoblotting. HEK293FT cells were from Thermo Fisher Scientific. *ARID1A-*knockout HCT116 (homozygous truncating mutations, Q456*/Q456*) and *ARID1A*-wildtype HCT116 colorectal carcinoma cells were from Horizon Discovery.

### Mass Spectrometry Analysis of the Immunopurified ARID1A Complex

The immunoprecipitated ARID1A protein complex was separated by gel electrophoresis, followed by peptide analysis of the digested gel bands by C18 reversed-phase chromatography using an UltiMate 3000 RSLCnano System (Thermo Scientific) equipped with an Acclaim PepMap C18 column (Thermo Scientific) and connected to a TriVersa NanoMate nanoelectrospray source (Advion) and a linear ion trap LTQ XL mass spectrometer (Thermo Scientific). Protein identification was performed using Mascot search engine v. 2.5.1 (Matrix Science) against the NCBI Homo sapiens database. Scaffold software v. 4.5.1 (Proteome Software) was used to validate the MS/MS peptide and protein identification based on 95% peptide and 99% protein probabilities, respectively.

### Functional similarities analysis

Gene Ontology (GO) enrichment analysis by R packages was conducted as previously described^22^. Briefly, based on the semantic similarities of GO terms used for gene annotation, protein inside the interactome were ranked by the average functional similarities between the protein and its interaction partners. Functional similarity, which is defined as the geometric mean of their semantic similarities in molecular function (MF), cellular component (CC) and biological process(BP) aspect of GO, was designed for measuring the strength of the relationship between each protein and its partners by considering function and location of proteins. The distributions of functional similarities were demonstrated in supplementary Fig. S1. Proteins, which showed strong relationship in function and location among the proteins within the interactome, were essential for the interactome to exert their functions. The average of functional similarities was used to rank protein in the ARID1A interactome. A cutoff value of 0.5 was chosen. The source code is available upon request.

### Immunoblot Analysis and Co-Immunoprecipitation Assay

For immunoblotting experiments, cells were lysed in BC200 lysis buffer [20 mM Tris-HCl (pH 7.5), 200 mM NaCl, 1 mM EDTA, 0.2% Nonidet P-40 (NP-40), freshly added complete protease inhibitor cocktail (Roche), and PhosSTOP phosphatase inhibitor (Roche)] or in boiling SDS lysis buffer [1% SDS and 10 mM Tris-Cl (pH 7.5)], as previously described^23^. Protein concentrations were quantified using a modified Lowry assay, and equal protein amounts were loaded onto a 10% SDS-PAGE gel and separated by gel electrophoresis, followed by transfer to a nitrocellulose membrane.

The membrane was stained with Ponceau S to confirm equal protein loading and then blocked in 2% BSA in Tris-buffered saline with 0.1% Tween-20 (TBST). Primary antibody incubation was done for 2 h at room temperature or, for phospho-specific antibodies, overnight at 4°C. An HRP-conjugated secondary antibody was used, followed by detection using enhanced chemiluminescence substrate (Pierce).

Autoradiograph images were scanned and saved as unmodified Tiff images, and densitometry analysis was done with ImageJ (NIH) software.

For co-immunoprecipitation, cells were lysed in BC150 lysis buffer [20 mM Tris-HCl (pH 7.5), 150 mM NaCl, 10% glycerol, 1 mM EDTA, 0.2% NP-40, and freshly added complete protease inhibitor cocktail (Roche)]. Cell lysates were incubated with primary antibody or IgG control antibody at 4°C overnight, followed by incubation with Protein A Agarose beads (Sigma-Aldrich) for 1 h at 4°C. After five washes with lysis buffer, the bound proteins were eluted from the beads in 2x Laemmli SDS sample loading buffer at 95°C for 5 min and then loaded onto a 10% SDS-PAGE gel. Proteins were transferred to nitrocellulose membranes, and immunoblotting was performed as described above.

### Glutathione S Transferase (GST) Protein-Protein Interaction Assay

Recombinant proteins, GST, GST-tagged CAD, and GST-tagged CAD fragments, shown in Fig. 2a, were purified as previously described^24^. HEK293T cells were transfected with pCIN4-HA-ARID1A using Lipofectamine 3000 (Invitrogen). Forty-eight hours later, the cells were lysed in lysis buffer [50 mM Tris-HCl (pH 8), 5 mM EDTA, 150 mM NaCl, 0.5% NP-40, and freshly added 1 mM DTT and complete protease inhibitor cocktail (Roche)]. The HEK293T lysates were then incubated for 2 h at 4°C with 25 μg Glutathione Sepharose 4B beads (Amersham) bound to GST-CAD or its fragments. After washing in lysis buffer, bound proteins were eluted from the beads with 2x Laemmli SDS sample loading buffer at 95°C for 5 min, loaded onto a 10% SDS-PAGE gel, and then transferred to a nitrocellulose membrane and immunoblotted using the indicated primary antibodies.

To evaluate domain of ARID1A that potentially interacts with GST-ATCase, HEK293T cells were transfected with pCDNA6-V5/His.b (empty vector), pcDNA6-ARID1A 1-1758 (N-terminus), pcDNA6-ARID1A 1759-2285 (C-terminus), or pcDNA6-ARID1A (full-length) expression plasmid using Lipofectamine 3000 (Invitrogen). Forty-eight hours later, the cells were lysed as described above. Lysates were then incubated for 2 h at 4°C with 25 μg Glutathione Sepharose 4B beads (Amersham) bound to GST or GST-ATCase. After washing in lysis buffer, bound proteins were eluted from the beads in 2x Laemmli SDS sample loading buffer at 95°C for 5 min and then loaded onto an SDS-PAGE gel, followed by transfer to a nitrocellulose membrane and immunoblotting using the indicated primary antibodies.

### Short Hairpin RNA (shRNA)-Mediated Knockdown and Expression of *ARID1A* in ovarian and endometrial carcinoma cells

shRNA lentiviral supernatants were produced using standard protocols, as previously described^11^. Vectors were transfected into HEK293FT cells using X-tremeGENE 9 DNA Transfection Reagent (Roche). Retroviral supernatants isolated at 48 h were diluted 1:1 in culture medium and used to infect the *ARID1A*-wildtype ES2 and KLE cell lines. Stably transfected subclones were expanded using puromycin (Dot Scientific) drug selection. For expression of ARID1A in the *ARID1A*-mutant HEC-1-A cell line, pCIN4-HA-ARID1A was transfected using X-tremeGENE 9 DNA Transfection Reagent (Roche), followed by selection with G418 (Gibco). For expression of ARID1A in the *ARID1A*-mutant SKOV3 and OVISE cell lines, V5/His-tagged pLenti-puro-LacZ and pLenti-puro-ARID1A were transfected using lentivirus. Lentivirus was produced using HEK293FT cells with the second-generation packaging system pSPAX2 (Addgene plasmid) and pMD2.G (Addgene plasmid). Stably transfected subclones were expanded using puromycin and blasticidin (Gibco) drug selection.

### Targeted exon sequencing for *ARID1A*

*ARID1A* mutations in SKOV3, A2780, HEC-1-A and OVISE cells were verified by Sanger sequencing using a Applied Biosystems 3730xL DNA Analyzer (Thermo Fisher Scientific, Inc.) and specific primers targeting exons 1, 2, 3, 18 and 20 (Supplementary Table S9) of the *ARID1A* (ENST00000324856) CDS region were used according to Jones, S. *et al*^2^. Briefly, Genomic DNA was extracted using QIAamp UCP DNA Micro Kit (Qiagen). PCR amplification with targeted primers was conducted using a touchdown PCR protocol (1 cycle of 96°C for 2 min; 3 cycles of 96°C for 10 sec, 64°C for 10 sec, 70°C for 30 sec; 3 cycles of 96°C for 10 sec, 61°C for 10 sec, 70°C for 30 sec; 3 cycles of 96°C for 10 sec, 58°C for 10 sec, 70°C for 30 sec; 41 cycles of 96°C for 10 sec, 57°C for 10 sec, 70°C for 30 sec; 1 cycle of 70°C for 5 min). PCR products were purified using QIAquick PCR Purification kit (Qiagen). PCR products were followed by Sanger sequencing. Mutations in *ARID1A* in those verified cell lines by Sanger sequencing was in Supplementary Table S10.

### Knockout of ARID1A in ES2 cell line

The CRISPR-Cas9 system was used to according to Ran *et al*^25^. The CRISPR/Cas9 vector, pSpCas9(BB)-2A-Puro (PX459) V2.0 (ID: 62988), was obtained from Addgene (MA, USA). The target site used in this study was 5′-CACCG<u>AGG</u>GAAGCGCTGCTGGGAAT-3′ that contains a part of the *ARID1A* sequence. The PAM sequence is underlined. The target sequence was inserted into the cloning site of the pSpCas9(BB)-2A-Puro (PX459) V2.0 vector. The cloned plasmid was transfected into ES2 cells using Lipofectamine 3000. ES2 cells were selected in the medium containing puromycin (1 μg/ml) 72 h after transfection and screening for single clones in about three weeks. *ARID1A*-null clones were identified by Western blotting and confirmed by Sanger sequencing the targeted genome region by PCR amplification with primers 5’-GTAAAACGACGGCCAGTTGCACGTTAGAGAACCACTCTG-3’ and 5’-AACAGCTATGACCATGACAACCAGCAAAGTCCTCACC-3’.

### ^14^C aspartate incorporation into RNA and DNA

Cells were plated in 60-mm dishes 48 h prior to the experiment (cells grew to 85-90% confluent when the experiment started). Fresh medium was added to the subconfluent cells, and 5 μCi L-[U-^14^C]aspartic acid (0.1 mCi/mL, PerkinElmer) was added to each plate. After 6 h of incubation at 37°C, the cells were lysed, and RNA and DNA were prepared following the manufacturer’s manual for the AllPrep DNA/RNA Mini Kit (Qiagen). The amounts of RNA and DNA were quantified, and the radioactivity in each sample was determined by liquid scintillation counting. [^14^C]Aspartate incorporation into RNA or DNA was respectively normalized to the amount of RNA or DNA and expressed as cpm/μg RNA or cpm/μg DNA.

### Uridine Triphosphate (UTP) assay

UTP was examined with an enzyme immuno-based plate-reader assay according to the manufacturer’s recommendations (Aviva Systems Biology). Briefly, cells were cultured in medium with or without drug treatment. For metabolite extraction, the medium was aspirated, and the same number of cells were collected by trypsinizing and counting. The cells were lysed by ultra-sonication (Qsonica Q125; Time: 2 min, Pulse: 15 s on/15 s off, Amplitude: 50%). The insoluble material in lysates was pelleted by centrifugation at 12,000 rpm for 10 min at 4°C. The metabolite-containing supernatants were assessed using the UTP ELISA Kit. The plate was read at 450 nm with a standard microplate reader. The UTP level was calculated using the formula (Relative OD_450_) = (Well OD_450_) – (Mean Blank Well OD_450_).

### Cytotoxicity assay

Cells were seeded in 96-well plates, with 2000 cells in 100 μL/well, and cultured for 24 h. The cells were treated with serial dilutions of the indicated drugs or without drug for an additional 72 h. The cell number was determined using the sulforhodamine B (SRB) assay, as previously described^26^. Briefly, cells were fixed with 20% trichloroacetic acid (TCA), air-dried, and stained with 0.4% SRB dissolved in 1% acetic acid. After washing, protein-bound dye was solubilized with 10 mM unbuffered Tris-base solution (pH 10.5) and detected at 510 nm using a microplate reader. Calculated the percentage of cell-growth using the following formula: % cell growth = Absorbance sample/Absorbance untreated × 100

Using CalcuSyn software (Biosoft), dose-effect curves were generated, and the drug concentrations corresponding to a 50% decrease in cell number (IC50) were determined.

### Immunofluorescence

Cells plated on coverslips were kept in medium. For detecting proteins, coverslips were fixed with 2% paraformaldehyde for 10 min, permeabilized with 0.5% Triton X-100 for 10 min, and blocked with 1% BSA in 20 mmol/L Tris-HCl (pH 7.5) for 20 min. The coverslips were then incubated with primary antibody (anti-ARID1A, 1:500; anti-CAD, 1:50; or P-CAD Ser1859, 1:50) overnight at 4°C and with secondary antibody for 2 h at room temperature. For multicolor staining, an additional blocking step was conducted after the first secondary antibody staining. DAPI was used to label the nucleus. Images of cells were acquired using a BZ-X710 fluorescence microscope (KEYENCE) and analyzed using ImageJ (NIH).

### Immunohistochemistry (IHC)

IHC was performed by incubating FFPE tissue section slides in antigen retrieval solution [0.01 M sodium citrate (pH 6.0)] for 20 min in a pressure cooker. The slides were blocked in blocking solution (5% goat serum and 2% BSA in TBS) for 30 min and then incubated with primary antibodies (anti-ARID1A, 1:500; anti-CAD, 1:50) overnight at 4°C. The slides were then incubated with the secondary antibody from the EnVision GI2 Doublestain System (DAKO) for 10 min at room temperature, followed by DAB staining for visualization. The slides were counterstained with hematoxylin and bluing in PBS, dehydrated in graded alcohol, cleared in xylene, and coverslipped in Permount (Fisher Scientific). Images were visualized using a BZ-X710 fluorescence microscope (KEYENCE) and analyzed using ImageJ (NIH).

### Animal model

#### Ethics statement

All animal procedures were conducted in accordance with a protocol approved by the Institutional Animal Care and Use Committee at Yale University (Protocol number: 2017-20139).

Five mouse experiments were performed: (i) to assess the effect of teriflunomide on ES2 cells *in vivo*, (ii) To assess the effect of teriflunomide on SKOV3 cells *in vivo*, (iii) to assess the effect of teriflunomide on patient-derived xenograft model, (iv) to assess the combination effect of teriflunomide and AZD6738 on ES2-shARID1A cells *in vivo* and (v) to assess the combination effect of teriflunomide and AZD6738 on patient-derived xenograft model. The i, ii, and iv experiments were experiments using subcutaneous cell-xenograft models generated by injecting cells into the flanks of 6-week-old female athymic NCr-nu/nu mice (Charles River Laboratories). The iii and v experiments were performed using patient-derived xenografts (PDXs). *Prkdc*^*em26Cd52*^*Il2rg*^*em26Cd22*^/NjuCrl (NCG) mice were purchased from the Charles River. PDXs were generated by sectioning of Cryo ovarian tumor tissue (Champion Oncology) and engrafting tumor chunks (5 × 5 × 5 mm) pieces subcutaneously to the 6- to 8-week-old female mice. Once the PDX tumor reached approximately 1,000 mm^3^, it was harvested and transplanted for expansion in next generations, which were used for *in vivo* studies. Animal-human dose translation was calculated as previously described^27^. Tumor volumes were measured every other day by caliper to determine tumor volume using the formula (length/2) × (width^2^). Animal weights were recorded, and mice were observed for any toxicities. The experiment was terminated when the mean tumor volume of the vehicle group reached 1000 mm^3^, and tumor xenografts were excised at the time of euthanasia. Representative samples were flash-frozen in liquid nitrogen for subsequent protein expression analysis by immunoblotting, as well as being formalin-fixed paraffin-embedded (FFPE) for subsequent hematoxylin and eosin staining and immunohistochemical analysis. FFPE sections were reviewed by the study pathologist (P.H.), and cellular necrosis was quantified as % cross-sectional area of the bisected tumor xenografts.

(i) To assess the effect of teriflunomide on ES2 cells, xenograft models were generated by injecting ES2-shCon or ES2-shARID1A cells (1 × 10^6^ cells) subcutaneously. When the mean tumor volume reached approximately 100 mm^3^, the animals were randomized into treatment groups; mice with xenograft volume <20 mm^3^ or >160 mm^3^ were excluded. Teriflunomide was solubilized in DMSO and diluted to 0.5 mg/mL with PBS. Mice were treated with teriflunomide (4 mg/kg) or vehicle intraperitoneally every other day, as shown in Fig. 4d.

(ii) To assess the effect of teriflunomide on SKOV3 cells, xenograft models were generated by injecting SKOV3 cells [2 × 10^6^ cells mixed 1:1 (v/v) with Matrigel (BD Biosciences)] subcutaneously. When the mean tumor volume reached approximately 100 mm^3^, the animals were randomized into treatment groups. Teriflunomide treatment was used the same way as in the ES2 xenograft models. The survival curve is shown in Fig. 4h; the terminal tumor volume was 1000 mm^3^. (iii) To assess the effect of teriflunomide on patient-derived xenografts (PDXs), the animals were randomized into treatment groups once the mean tumor volume reached approximately 100 mm^3^. Teriflunomide treatment was used the same way as in the ES2 xenograft models. Tumor size was monitored every four days by a caliper, as shown in Fig. 4i. The terminal tumor volume was 1000 mm^3^.

(iv) To assess the combination effect of teriflunomide and AZD6738 on ES2-shARID1A cells *in vivo*, xenograft models were generated by injecting ES2-shARID1A cells [2 × 10^6^ cells mixed 1:1 (v/v) with Matrigel (BD Biosciences)] subcutaneously. Treatment with was initiated 24 h after tumor injection, the animals were randomized into treatment groups. Teriflunomide treatment was used the same way as in the ES2 xenograft models. AZD6738 was solubilized in DMSO and diluted to 0.5 mg/mL with 10% 2-hydroxypropyl-b-cyclodextrin. Treatment with AZD6738 (25 mg/kg) or vehicle was performed daily by oral gavage, as shown in Fig. 5l.

(v) To assess the combination effect of teriflunomide and AZD6738 on patient-derived xenografts (PDXs), ARID1A deficient ovarian tumor tissues (Champion Oncology, CTG-2213) were used in the study. The animals were randomized into treatment groups once the mean tumor volume reached approximately 100 mm^3^. Teriflunomide and AZD6738 treatment was used the same way as in the ES2 xenograft models. Tumor size was monitored every four days by a caliper, as shown in Fig. 5m. The terminal tumor volume was 500 mm^3^ in vehicle group.

### Statistics

An independent-samples t-test was applied when two groups of data were compared. Multiple-group comparisons were done using one-way analysis of variance (ANOVA) with Tukey’s post-test. Statistical analyses and graphing were performed using SPSS 22 (IBM) and Prism 7 (GraphPad). *P*-values less than 0.05 were considered significant.

## Acknowledgements

Funding for this research was provided by the Department of Defense Ovarian Cancer Research Program Award W81XWH-16-1-0196 and the Reproductive Scientist Development Program Seed Grant to G. S. Huang. We thank Dr. Jennifer T. Aguilan and the comprehensive mass spectrometry services provided by the Proteomics Shared Resource of the Albert Einstein College of Medicine, supported by the Cancer Center Support Grant (NCI P30 CA013330). We thank the Shared Resources of the Yale Cancer Center, supported by the Cancer Center Support Grant (NCI P30 P30 CA016359), and the YCC Scientific Publication Program supported by The Sands Family Foundation. We thank Dr. Ie-Ming Shih for providing ARID1A vectors and short hairpin RNA vectors used in this study. We acknowledge the Yale Genome Editing Center for providing custom CRISPR/Cas constructs. We thank Dr. Susan Band Horwitz for helpful discussions.

## Authors’ Contributions

**Conception and design:** G.S. Huang, Z. Li, S. Mi

**Development of methodology:** Z. Li, S. Mi, C-P.H. Yang, G.S. Huang

**Acquisition of data (provided animals, acquired and managed patients, provided facilities, etc.):** Z. Li, S. Mi, O.I. Osagie, J. Ji, C-P.H. Yang, M. Schwartz, P. Hui, G.S. Huang

**Analysis and interpretation of data (e.g., statistical analysis, biostatistics, computational analysis):** Z. Li, S. Mi, O.I. Osagie, J. Ji, C-P.H. Yang, M. Schwartz, P. Hui, G.S. Huang

**Writing, review, and/or revision of the manuscript:** Z. Li, S. Mi, O.I. Osagie, J. Ji, C-P.H. Yang, M. Schwartz, P. Hui, G.S. Huang

**Study supervision:** G.S. Huang

## Current Affiliations

Shijun Mi, Laboratory of Personalized Genomic Medicine, Department of Pathology & Cell Biology, Columbia University Medical Center, New York. Jing Ji, Department of Obstetrics & Gynecology, the First Affiliated Hospital of Medical School, Xi’an Jiaotong University, Xi’an 710061, The People’s Republic of China. Oloruntoba I. Osagie, Queen’s University Belfast, Northern Ireland, United Kingdom. Melissa Schwartz, Department of Obstetrics, Gynecology and Women’s Health, Division of Gynecologic Oncology, Saint Louis University School of Medicine, St. Louis, Missouri.

## Corresponding Author

Correspondence to G.S. Huang.

## REFERENCES

1 Wiegand, K. C. et al. ARID1A mutations in endometriosis-associated ovarian carcinomas. New England Journal of Medicine 363, 1532–1543 (2010).

2 Jones, S. et al. Frequent mutations of chromatin remodeling gene ARID1A in ovarian clear cell carcinoma. Science 330, 228–231 (2010).

3 Wu, R.-C., Wang, T.-L. & Shih, I.-M. The emerging roles of ARID1A in tumor suppression. Cancer biology & therapy 15, 655–664 (2014).

4 Maeda, D. et al. Clinicopathological significance of loss of ARID1A immunoreactivity in ovarian clear cell carcinoma. International journal of molecular sciences 11, 5120–5128 (2010).

5 Wilson, B. G. & Roberts, C. W. SWI/SNF nucleosome remodellers and cancer. Nature Reviews Cancer 11, 481 (2011).

6 Li, J. et al. Epigenetic driver mutations in ARID1A shape cancer immune phenotype and immunotherapy. The Journal of Clinical Investigation (2020).

7 Katagiri, A. et al. Loss of ARID1A expression is related to shorter progression-free survival and chemoresistance in ovarian clear cell carcinoma. Modern Pathology 25, 282–288 (2012).

8 Forbes, S. A. et al. COSMIC: somatic cancer genetics at high-resolution. Nucleic Acids Research 45, D777–D783 (2016).

9 Anglesio, M. S. et al. Type-specific cell line models for type-specific ovarian cancer research. PloS one 8, e72162 (2013).

10 Dykhuizen, E. C. et al. BAF complexes facilitate decatenation of DNA by topoisomerase IIα. Nature 497, 624 (2013).

11 Guan, B., Wang, T.-L. & Shih, I.-M. ARID1A, a factor that promotes formation of SWI/SNF-mediated chromatin remodeling, is a tumor suppressor in gynecologic cancers. Cancer research 71, 6718–6727 (2011).

12 Ben-Sahra, I., Howell, J. J., Asara, J. M. & Manning, B. D. Stimulation of de novo pyrimidine synthesis by growth signaling through mTOR and S6K1. Science 339, 1323–1328 (2013).

13 Robitaille, A. M. et al. Quantitative phosphoproteomics reveal mTORC1 activates de novo pyrimidine synthesis. Science 339, 1320–1323 (2013).

14 Barretina, J. et al. The Cancer Cell Line Encyclopedia enables predictive modelling of anticancer drug sensitivity. Nature 483, 603 (2012).

15 Ruiz-Ramos, A., Velázquez-Campoy, A., Grande-García, A., Moreno-Morcillo, M. & Ramón-Maiques, S. Structure and functional characterization of human aspartate transcarbamoylase, the target of the anti-tumoral drug PALA. Structure 24, 1081–1094 (2016).

16 O’Connor, P. et al. Randomized trial of oral teriflunomide for relapsing multiple sclerosis. New England Journal of Medicine 365, 1293–1303 (2011).

17 Williamson, C. T. et al. ATR inhibitors as a synthetic lethal therapy for tumours deficient in ARID1A. Nature Communications 7 (2016).

18 Chou, T.-C. Drug combination studies and their synergy quantification using the Chou-Talalay method. Cancer research 70, 440–446 (2010).

19 Wild, J., Loughrey-Chen, S. & Corder, T. In the presence of CTP, UTP becomes an allosteric inhibitor of aspartate transcarbamoylase. Proceedings of the National Academy of Sciences 86, 46–50 (1989).

20 Rabinowitz, J. D. et al. Dissecting enzyme regulation by multiple allosteric effectors: nucleotide regulation of aspartate transcarbamoylase. Biochemistry 47, 5881–5888 (2008).

21 Rees, S. et al. Bicistronic vector for the creation of stable mammalian cell lines that predisposes all antibiotic-resistant cells to express recombinant protein. Biotechniques 20, 102-104, 106, 108–110 (1996).

22 Han, Y. et al. Proteomic investigation of the interactome of FMNL1 in hematopoietic cells unveils a role in calcium-dependent membrane plasticity. Journal of proteomics 78, 72–82 (2013).

23 Yang, C.-P. H. & Horwitz, S. B. Taxol mediates serine phosphorylation of the 66-kDa Shc isoform. Cancer Research 60, 5171–5178 (2000).

24 Keller, D. M., Zeng, S. X. & Lu, H. in p53 Protocols. 121–133 (Springer, Totowa, NJ, 2003).

25 Ran, F. A. et al. Genome engineering using the CRISPR-Cas9 system. Nature protocols 8, 2281 (2013).

26 Huang, G. S. et al. Insulin-like growth factor 2 expression modulates Taxol resistance and is a candidate biomarker for reduced disease-free survival in ovarian cancer. Clinical Cancer Research 16, 2999–3010 (2010).

27 Reagan-Shaw, S., Nihal, M. & Ahmad, N. Dose translation from animal to human studies revisited. The FASEB Journal 22, 659–661 (2008).

