## Supplemental tables and figures for "Vulnerability of ARID1A deficient cancer cells to pyrimidine synthesis blockade"

### TITLE

Table S1. Analysis of the combination of AZD6738 with FDA-approved teriflunomide in ES2 cells.

| Cell Type And Drug | CI Values at | | | Dm | r | $CI_{wt}$ |
|---|---|---|---|---|---|---|
| | ED50 | ED75 | ED90 | | | |
| ES2-Teriflunomide | N/A | N/A | N/A | 26.36 ± 2.18 | 0.98 ± 0.009 | N/A |
| SCR-Teriflunomide | N/A | N/A | N/A | 28.15 ± 1.59 | 0.98 ± 0.003 | N/A |
| shARID1A (a)-Teriflunomide | N/A | N/A | N/A | 17.86 ± 1.32*** | 0.91 ± 0.052 | N/A |
| shARID1A (b)-Teriflunomide | N/A | N/A | N/A | 16.76 ± 0.92*** | 0.99 ± 0.003 | N/A |
| ES2-AZD6738 | N/A | N/A | N/A | 1.23 ± 0.16 | 0.92 ± 0.020 | N/A |
| SCR-AZD6738 | N/A | N/A | N/A | 1.18 ± 0.17 | 0.92 ± 0.026 | N/A |
| shARID1A (a)-AZD6738 | N/A | N/A | N/A | 0.20 ± 0.01*** | 0.88 ± 0.022 | N/A |
| shARID1A (b)-AZD6738 | N/A | N/A | N/A | 0.24 ± 0.03*** | 0.85± 0.016 | N/A |
| ES2-Combo (10:1) | 2.51 ± 0.251 | 0.68 ± 0.076 | 0.22 ± 0.030 | 7.57 ± 0.50 | 0.98 ± 0.002 | 0.76 ± 0.049 |
| SCR-Combo (10:1) | 2.21 ± 0.154 | 0.55 ± 0.054 | 0.17 ± 0.050 | 7.85 ± 1.05 | 0.99 ± 0.010 | 0.64 ± 0.020 |
| shARID1A (a)-Combo (10:1) | 0.91 ± 0.099 | 0.53 ± 0.007 | 0.32 ± 0.028 | 4.57 ± 0.18** | 0.98 ± 0.006 | 0.49 ± 0.005 |
| shARID1A (b)-Combo (10:1) | 0.95 ± 0.152 | 0.54 ± 0.091 | 0.31 ± 0.065 | 4.62 ± 0.19** | 0.98 ± 0.004 | 0.49 ± 0.083 |

Table S1. CI values were determined for the indicated two-drug combinations at 50% (ED50), 75% (ED75), and 90% (ED90) inhibition of ES2 cell proliferation using the CalcuSyn program. The $CI_{wt}$ showed the overall synergic effect of combination tests. $CI_{wt}$ < 0.8, = 0.8 to 1.2, and > 1.2 indicates synergism, an additive effect, and antagonism, respectively. The $CI_{wt}$ value was assigned as follows: (ED50 + 2ED75 + 3ED90)/6. Datasets from at least three independent experiments were combined, and means ± SD were calculated for each combination. SCR, ES2-shScrambled. N/A, not applicable. Statistical analysis was carried out by one-way ANOVA with Bonferroni's post-hoc test, and differences were considered significant at $** P < 0.01$ and $*** P < 0.001$ as compared to the ES2 parental group with the same drug treatment.

Table S2. Analysis of the combination of AZD6738 with teriflunomide in A2780 cells.

| Drug | CI Values at | | | Dm | m | R | $CI_{wt}$ |
|---|---|---|---|---|---|---|---|
| | ED50 | ED75 | ED90 | | | | |
| Teriflunomide | N/A | N/A | N/A | 18.91 ± 1.39 | 0.82 ± 0.13 | 0.99 ± 0.001 | N/A |
| AZD6738 | N/A | N/A | N/A | 5.54 ± 1.06 | 1.19 ± 0.18 | 0.99 ± 0.003 | N/A |
| Combo (10:1) | 0.74 ± 0.03 | 0.47 ± 0.05 | 0.31 ± 0.08 | 10.40 ± 0.82 | 1.48 ± 0.35 | 0.99 ± 0.003 | 0.44 ± 0.06 |

Table S2. CI values were determined for the indicated two-drug combinations at 50% (ED50), 75% (ED75), and 90% (ED90) inhibition of A2780 cell proliferation using the CalcuSyn program. The $CI_{wt}$ showed the overall synergic effect of combination tests. $CI_{wt}$ < 0.8, = 0.8 to 1.2, and > 1.2 indicate synergism, an additive effect, and antagonism, respectively. The $CI_{wt}$ value was assigned as follows: (ED50 + 2ED75 + 3ED90)/6. Datasets from at least three independent experiments were combined, and means ± SD were calculated for each combination.

Table S3. Analysis of the combination of AZD6738 with teriflunomide in JHOC-5 cells.

| Drug | CI Values at | | | Dm | m | R | $CI_{wt}$ |
|---|---|---|---|---|---|---|---|
| | ED50 | ED75 | ED90 | | | | |
| Teriflunomide | N/A | N/A | N/A | 25.20 ± 1.85 | 1.10 ± 0.03 | 0.98 ± 0.006 | N/A |
| AZD6738 | N/A | N/A | N/A | 2.91 ± 0.76 | 1.20 ± 0.12 | 0.99 ± 0.012 | N/A |
| Combo (10:1) | 1.03 ± 0.03 | 0.82 ± 0.05 | 0.66 ± 0.09 | 13.64 ± 1.42 | 1.52 ± 0.22 | 0.99 ± 0.008 | 0.78 ± 0.06 |

Table S3. CI values were determined for the indicated two-drug combinations at 50% (ED50), 75% (ED75), and 90% (ED90) inhibition of JHOC-5 cell proliferation using the CalcuSyn program. The $CI_{wt}$ showed the overall synergic effect of combination tests. $CI_{wt}$ < 0.8, = 0.8 to 1.2, and > 1.2 indicates synergism, an additive effect, and antagonism, respectively. The $CI_{wt}$ value was assigned as follows: (ED50 + 2ED75 + 3ED90)/6. Datasets from at least three independent experiments were combined, and means ± SD were calculated for each combination.

Table S4. Analysis of the combination of AZD6738 with teriflunomide in HEC-1-A cells.

| Drug | CI Values at | | | Dm | m | R | $CI_{wt}$ |
|---|---|---|---|---|---|---|---|
| | ED50 | ED75 | ED90 | | | | |
| Teriflunomide | N/A | N/A | N/A | 50.49 ± 4.99 | 0.85 ± 0.26 | 0.99 ± 0.005 | N/A |
| AZD6738 | N/A | N/A | N/A | 4.47 ± 0.45 | 1.25 ± 0.20 | 0.99 ± 0.003 | N/A |
| Combo (10:1) | 0.81 ± 0.10 | 0.67 ± 0.24 | 0.61 ± 0.30 | 19.06 ± 0.61 | 1.36 ± 0.38 | 0.99 ± 0.004 | 0.66 ± 0.25 |

Table S4. CI values were determined for the indicated two-drug combinations at 50% (ED50), 75% (ED75), and 90% (ED90) inhibition of HEC-1A cell proliferation using the CalcuSyn program. The $CI_{wt}$ showed the overall synergic effect of combination tests. $CI_{wt}$ < 0.8, = 0.8 to 1.2, and > 1.2 indicates synergism, an additive effect, and antagonism, respectively. The $CI_{wt}$ value was assigned as follows: (ED50 + 2ED75 + 3ED90)/6. Datasets from at least three independent experiments were combined, and means ± SD were calculated for each combination.

Table S5. Analysis of the combination of VX-970 with teriflunomide in ES2 cells.

| Cell Type And Drug | CI Values at | | | Dm | R | $CI_{wt}$ |
|---|---|---|---|---|---|---|
| | ED50 | ED75 | ED90 | | | |
| ES2-Teri (μM) | N/A | N/A | N/A | 29.09 ± 0.54 | 0.97 ± 0.004 | N/A |
| SCR-Teri (μM) | N/A | N/A | N/A | 26.67 ± 1.26 | 0.97 ± 0.002 | N/A |
| shARID1A (a)-Teri (μM) | N/A | N/A | N/A | 15.57 ± 1.79*** | 0.99 ± 0.005 | N/A |
| shARID1A (b)-Teri (μM) | N/A | N/A | N/A | 15.46 ± 3.11*** | 0.98 ± 0.011 | N/A |
| ES2-VX970 (nM) | N/A | N/A | N/A | 165.22 ± 3.84 | 0.95 ± 0.004 | N/A |
| SCR-VX970 (nM) | N/A | N/A | N/A | 169.55 ± 9.24 | 0.96 ± 0.008 | N/A |
| shARID1A (a)-VX970 (nM) | N/A | N/A | N/A | 54.33 ± 1.31*** | 0.98 ± 0.005 | N/A |
| shARID1A (b)-VX970 (nM) | N/A | N/A | N/A | 63.32 ± 10.37*** | 0.98± 0.003 | N/A |
| ES2-Combo (200:1) | 0.63 ± 0.053 | 0.51 ± 0.015 | 0.42 ± 0.015 | 18.26 ± 1.72 | 0.98 ± 0.002 | 0.48 ± 0.026 |
| SCR-Combo (200:1) | 0.68 ± 0.078 | 0.53 ± 0.060 | 0.41 ± 0.060 | 18.03 ± 2.31 | 0.97 ± 0.010 | 0.50 ± 0.058 |
| shARID1A (a)-Combo (200:1) | 0.32 ± 0.020 | 0.30 ± 0.009 | 0.30 ± 0.009 | 4.92 ± 0.75 | 0.95 ± 0.006 | 0.31± 0.009 |
| shARID1A (b)-Combo (200:1) | 0.33 ± 0.023 | 0.31 ± 0.101 | 0.31 ± 0.101 | 5.08 ± 0.88*** | 0.93 ± 0.004 | 0.31 ± 0.119 |

Table S5. CI values were determined for the indicated two-drug combinations at 50% (ED50), 75% (ED75), and 90% (ED90) inhibition of ES2 cell proliferation using the CalcuSyn program. The $CI_{wt}$ showed the overall synergic effect of combination tests. $CI_{wt}$ < 0.8, = 0.8 to 1.2, and > 1.2 indicates synergism, an additive effect, and antagonism, respectively. The $CI_{wt}$ value was assigned as follows: (ED50 + 2ED75 + 3ED90)/6. Datasets from at least three independent experiments were combined, and means ± SD were calculated for each combination. Teri, teriflunomide. SCR, ES2-shScrambled. N/A, not applicable. Statistical analysis was carried out by one-way ANOVA with Bonferroni's post-hoc test, and differences were considered significant at $*** \ P < 0.001$ as compared to the ES2 parental group with the same drug treatment.

Table S6. Analysis of the combination of AZD6738 with FDA-approved teriflunomide in ARID1A-restoration OVISE cells.

| Cell Type And Drug | CI Values at | | | Dm | r | $CI_{wt}$ |
|---|---|---|---|---|---|---|
| | ED50 | ED75 | ED90 | | | |
| LacZ-Teri (μM) | N/A | N/A | N/A | 18.24 ± 1.19*** | 0.98 ± 0.025 | N/A |
| LacZ Dox-Teri (μM) | N/A | N/A | N/A | 19.30 ± 1.24*** | 0.99 ± 0.008 | N/A |
| ARID1A-Teri (μM) | N/A | N/A | N/A | 19.08 ± 1.07*** | 0.99 ± 0.001 | N/A |
| ARID1A Dox-Teri (μM) | N/A | N/A | N/A | 109.48 ± 3.18 | 0.98 ± 0.011 | N/A |
| LacZ-AZD6738 (μM) | N/A | N/A | N/A | 0.70 ± 0.03*** | 0.95 ± 0.030 | N/A |
| LacZ Dox-AZD6738 (μM) | N/A | N/A | N/A | 0.67 ± 0.11*** | 0.96 ± 0.021 | N/A |
| ARID1A-AZD6738 (μM) | N/A | N/A | N/A | 0.77 ± 0.16*** | 0.98 ± 0.012 | N/A |
| ARID1A Dox-AZD6738 (μM) | N/A | N/A | N/A | 6.99 ± 0.34 | 0.98± 0.014 | N/A |
| LacZ-Combo (10:1) | 1.30 ± 0.160 | 0.70 ± 0.048 | 0.40 ± 0.047 | 6.55 ± 1.03*** | 0.94 ± 0.013 | 0.65 ± 0.047 |
| LacZ Dox-Combo (10:1) | 1.16 ± 0.220 | 0.64 ± 0.030 | 0.39 ± 0.035 | 5.70 ± 0.81*** | 0.96 ± 0.002 | 0.60 ± 0.035 |
| ARID1A-Combo (10:1) | 1.18 ± 0.088 | 0.84 ± 0.182 | 0.65 ± 0.196 | 6.41 ± 0.48*** | 0.95 ± 0.013 | 0.80 ± 0.196 |
| ARID1A Dox-Combo (10:1) | 6.15 ± 0.254 | 3.21 ± 1.026 | 1.86 ± 0.934 | 262.45 ± 20.20 | 0.99 ± 0.004 | 3.02 ± 0.934 |

Table S6. CI values were determined for the indicated two-drug combinations at 50% (ED50), 75% (ED75), and 90% (ED90) inhibition of OVISE cell proliferation using the CalcuSyn program. The $CI_{wt}$ showed the overall synergic effect of combination tests. $CI_{wt}$ < 0.8, = 0.8 to 1.2, and > 1.2 indicates synergism, an additive effect, and antagonism, respectively. The $CI_{wt}$ value was assigned as follows: (ED50 + 2ED75 + 3ED90)/6. Datasets from at least three independent experiments were combined, and means ± SD were calculated for each combination. Teri, teriflunomide. N/A, not applicable. Statistical analysis was carried out by one-way ANOVA with Bonferroni's post-hoc test, and differences were considered significant at $*** P < 0.001$ as compared to the ARID1A restoration group induced by doxycycline with the same drug treatment.

Table S7. Analysis of the combination of AZD6738 with FDA-approved teriflunomide in ARID1A-restoration SKOV3 cells.

| | CI Values at | | | | | |
|---|---|---|---|---|---|---|
| **Cell Type And Drug** | **ED50** | **ED75** | **ED90** | **Dm** | **r** | $CI_{wt}$ |
| LacZ-Teri (µM) | N/A | N/A | N/A | 24.78 ± 2.45 | 0.95 ± 0.010 | N/A |
| LacZ-Dox-Teri (µM) | N/A | N/A | N/A | 23.46 ± 2.58 | 0.96 ± 0.018 | N/A |
| ARID1A-Teri (µM) | N/A | N/A | N/A | 23.93 ± 2.63 | 0.95 ± 0.021 | N/A |
| ARID1A-Dox-Teri (µM) | N/A | N/A | N/A | 73.41 ± 3.76*** | 0.98 ± 0.005 | N/A |
| LacZ-AZD6738 (µM) | N/A | N/A | N/A | 3.39 ± 0.13 | 0.97 ± 0.009 | N/A |
| LacZ-Dox-AZD6738 (µM) | N/A | N/A | N/A | 3.48 ± 0.70 | 0.96 ± 0.018 | N/A |
| ARID1A-AZD6738 (µM) | N/A | N/A | N/A | 3.40 ± 0.51 | 0.97 ± 0.010 | N/A |
| ARID1A-Dox-AZD6738 (µM) | N/A | N/A | N/A | 8.54 ± 0.58*** | 0.97 ± 0.028 | N/A |
| LacZ-Combos (5:1) | 0.65 ± 0.026 | 0.52 ± 0.068 | 0.42 ± 0.100 | 6.54 ± 0.41 | 0.96 ± 0.008 | 0.49 ± 0.075 |
| LacZ-Dox-Combos (5:1) | 0.70 ± 0.075 | 0.53 ± 0.051 | 0.42 ± 0.082 | 6.94 ± 0.33 | 0.94 ± 0.008 | 0.51 ± 0.051 |
| ARID1A-Combos (5:1) | 0.67 ± 0.075 | 0.56 ± 0.101 | 0.48 ± 0.124 | 6.73 ± 1.51 | 0.95 ± 0.026 | 0.54 ± 0.103 |
| ARID1A-Dox-Combos (5:1) | 2.27 ± 0.094 | 2.00 ± 0.327 | 1.81 ± 0.525 | 61.13 ± 3.06*** | 0.92 ± 0.004 | 1.95 ± 0.381 |

Table S7. CI values were determined for the indicated two-drug combinations at 50% (ED50), 75% (ED75), and 90% (ED90) inhibition of SKOV3 cell proliferation using the CalcuSyn program. The $CI_{wt}$ showed the overall synergic effect of combination tests. $CI_{wt}$ < 0.8, = 0.8 to 1.2, and > 1.2 indicates synergism, an additive effect, and antagonism, respectively. The $CI_{wt}$ value was assigned as follows: (ED50 + 2ED75 + 3ED90)/6. Datasets from at least three independent experiments were combined, and means ± SD were calculated for each combination. Teri, teriflunomide. N/A, not applicable. Statistical analysis was carried out by one-way ANOVA with Bonferroni's post-hoc test, and differences were considered significant at $*** P < 0.001$ as compared to the ARID1A restoration group induced by doxycycline with the same drug treatment.

Table S8. Analysis of the combination of AZD6738 with FDA-approved teriflunomide in *ARID1A*-knockout ES2 cells.

| Cell Type And Drug | CI Values at | | | Dm | r | $CI_{wt}$ |
|---|---|---|---|---|---|---|
| | ED50 | ED75 | ED90 | | | |
| ES2-Teri (μM) | N/A | N/A | N/A | 40.29 ± 2.49 | 0.98 ± 0.008 | N/A |
| ES2-vector-Teri (μM) | N/A | N/A | N/A | 43.73 ±3.01 | 0.97 ± 0.007 | N/A |
| ES2-sg1-Teri (μM) | N/A | N/A | N/A | 12.00 ± 0.75**** | 0.98 ± 0.012 | N/A |
| ES2-sg2-Teri (μM) | N/A | N/A | N/A | 12.50 ± 1.79*** | 0.98 ± 0.003 | N/A |
| ES2-AZD (μM) | N/A | N/A | N/A | 3.05 ± 0.57 | 0.93 ± 0.016 | N/A |
| ES2-vector-AZD (μM) | N/A | N/A | N/A | 3.10 ± 0.48 | 0.94 ± 0.012 | N/A |
| ES2-sg1-AZD (μM) | N/A | N/A | N/A | 1.63 ± 0.15* | 0.95 ± 0.0040 | N/A |
| ES2-sg2-AZD (μM) | N/A | N/A | N/A | 1.76 ± 0.22* | 0.93 ± 0.016 | N/A |
| ES2-Combos (20:1) | 0.95 ± 0.108 | 0.55 ± 0.026 | 0.33 ± 0.059 | 22.75 ± 0.23 | 0.99 ± 0.004 | 0.51 ± 0.026 |
| ES2-vector-Combos (20:1) | 0.92 ± 0.139 | 0.56 ± 0.070 | 0.35 ± 0.079 | 23.27 ± 2.63 | 0.96 ± 0.029 | 0.51 ± 0.062 |
| ES2-sg1-Combos (20:1) | 0.69 ± 0.037 | 0.55 ± 0.30 | 0.45 ± 0.020 | 6.10 ± 0.24*** | 0.99 ± 0.006 | 0.52 ± 0.026 |
| ES2-sg2-Combos (20:1) | 0.70 ± 0.100 | 0.56 ± 0.018 | 0.46 ± 0.037 | 6.36 ± 0.24*** | 0.99 ± 0.005 | 0.53 ± 0.011 |

Table S8. CI values were determined for the indicated two-drug combinations at 50% (ED50), 75% (ED75), and 90% (ED90) inhibition of SKOV3 cell proliferation using the CalcuSyn program. The $CI_{wt}$ showed the overall synergic effect of combination tests. $CI_{wt}$ < 0.8, = 0.8 to 1.2, and > 1.2 indicates synergism, an additive effect, and antagonism, respectively. The $CI_{wt}$ value was assigned as follows: (ED50 + 2ED75 + 3ED90)/6. Datasets from at least three independent experiments were combined, and means ± SD were calculated for each combination. Teri, teriflunomide. N/A, not applicable. Statistical analysis was carried out by one-way ANOVA with Bonferroni's post-hoc test, and differences were considered significant at $* P < 0.05$, $*** P < 0.001$ as compared to the PX459 empty vector group.

Table S9. PCR primers used for targeted PCR amplification and sequencing

| Name | Coding Exon Number | M13_F PCR primer sequence | Name | M13_R PCR primer sequence | Product length (bp) |
|---|---|---|---|---|---|
| **ARID1A_03** | 1 | GTAAAACGACGGCCAGTGGGAAAGGAGCTGCAGGA | ARID1A_4 | AACAGCTATGACCATGACCTCTCGGGGAGCTCAG | 502 |
| **ARID1A_09** | 2 | GTAAAACGACGGCCAGTTTGGAAGCCAAGGATACATTC | ARID1A_10 | AACAGCTATGACCATGAGGTTGGTCTCATTGCTCTTTC | 442 |
| **ARID1A_13** | 3 | GTAAAACGACGGCCAGTTGCACGTTAGAGAACCACTCTG | ARID1A_14 | AACAGCTATGACCATGACAACCAGCAAAGTCCTCACC | 495 |
| **ARID1A_43** | 18 | GTAAAACGACGGCCAGTGGAAGAAAGAGTGGTGGTTGC | ARID1A_44 | AACAGCTATGACCATGCCAAACTGGAATGGAAATTGG | 458 |
| **ARID1A_53** | 20 | GTAAAACGACGGCCAGTGTCTTGCTCTCGAAGTGGGTC | ARID1A_54 | AACAGCTATGACCATGGGAGAACCTTTGGGAAAGGAG | 599 |
| **ARID1A_61** | 20 | GTAAAACGACGGCCAGTCCTTGGTTACACTCGCCAAC | ARID1A_62 | AACAGCTATGACCATGCAGCCGTGATTCGTACAGAGTA | 596 |

Table S10. Mutations in *ARID1A* in human ovarian cell lines

| Gene Symbol | Coding Exon Number | SKOV3 CDS mutation | A2780 CDS mutation | HEC-1-A CDS mutation | OVISE CDS mutation |
|---|---|---|---|---|---|
| **ARID1A** | 1 | | | | c.606_607delTC (Deletion - Frameshift)<br>c.607_608insA (Insertion - Frameshift)<br>c.608_609insA (Insertion - Frameshift) |
| **ARID1A** | 2 | | | ~c1212 Q404H. Missense | |
| **ARID1A** | 3 | c.1756C>T (GRCh38, 1:26731557. p.Q586*. Nonsense) | | | |
| **ARID1A** | 18 | | Q1430* Nonsense (~c.4288) | | |
| **ARID1A** | 20 | | c.5161C>T (chr1:26978137. p.R1721*. Nonsense)) | G1761C. c.5281. Missense<br>c.5503C>T (chr1:26978479. Q1835*. Nonsense) | |
| **ARID1A** | 20 | | | Q2115* Nonsense (~chr1:26979621) | |

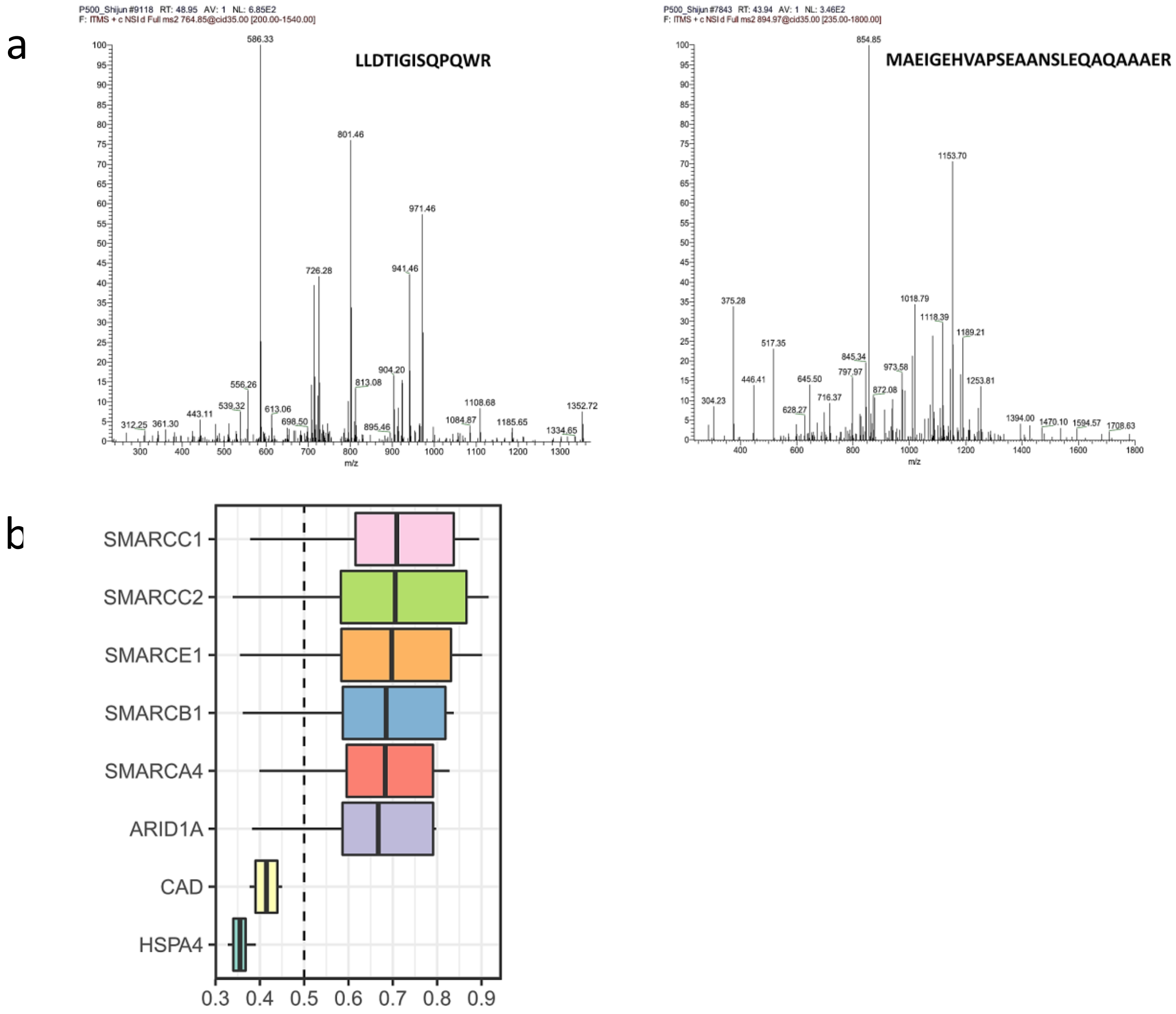


Figure S1. Representative fragmentation spectra of CAD peptides and functional similarities analysis of ARID1A interactome. **a**, representative fragmentation spectra of CAD peptides. **b**, summary of functional similarities of the ARID1A interactome in KLE cell. The distributions of functional similarities were summarized as boxplots. The boxes represent the middle 50% of the similarities; the upper and lower boundaries show the 75th and 25th percentile. The lines in the boxes indicate the mean of the functional similarities. Proteins with a higher average functional similarity (cutoff > 0.5) are considered as the known central proteins within the ARID1A interactome in KLE cell. The dashed line represents the cutoff value.

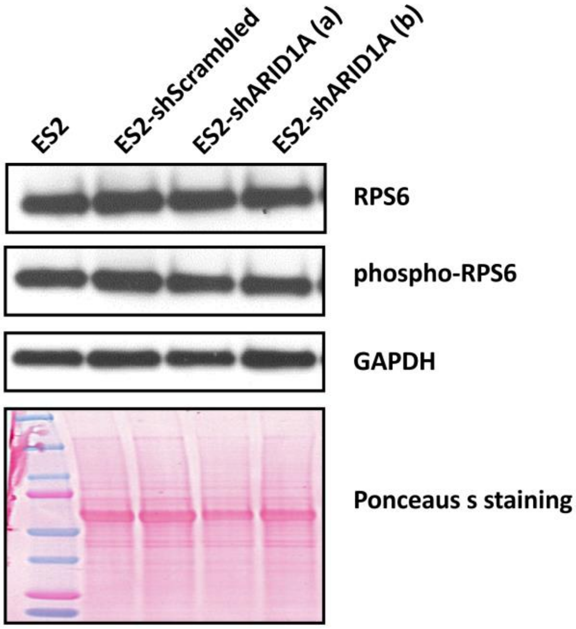


Figure S2. Unchanged RPS6 and phosphorylated RPS6 in *ARID1A* knockdown cells compared to control cells, shown in a representative immunoblot.

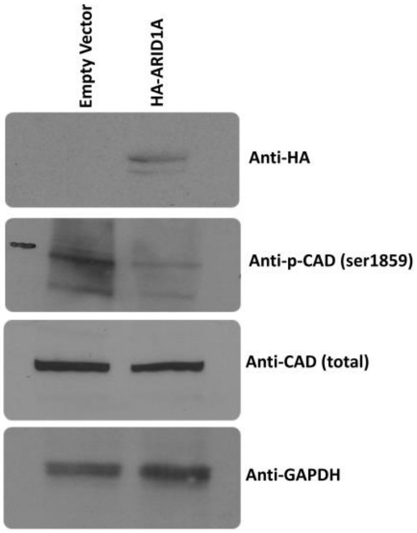


Figure S3. Restoration of ARID1A expression in the *ARID1A*-mutant cell line HEC-1-A leads to a reduction in phosphorylated and total CAD.

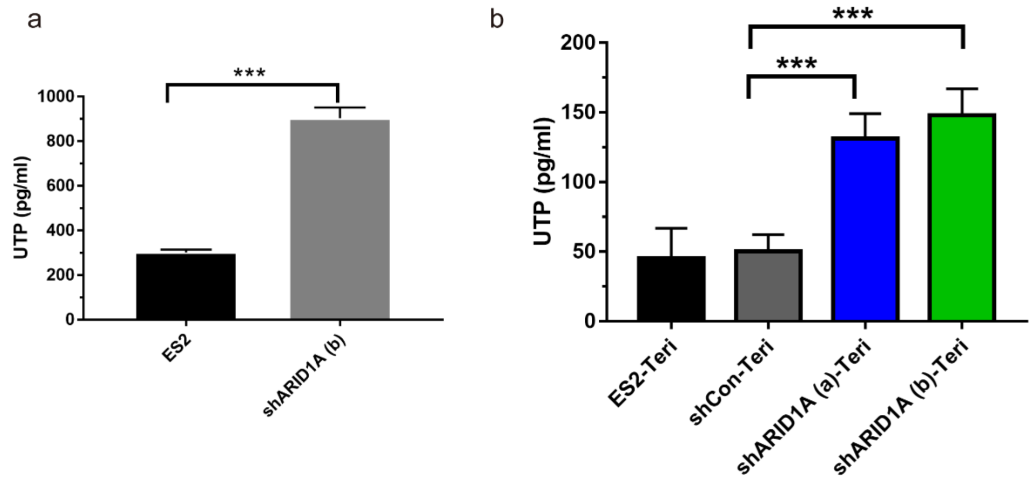


Figure S4. Increased UTP level shows in *ARID1A* knockdown ES2 cells. Teriflunomide decreases UTP abundance in both *ARID1A* wild-type and knockdown ES2 cells, but still maintaining higher pool in *ARID1A* knockdown cells.

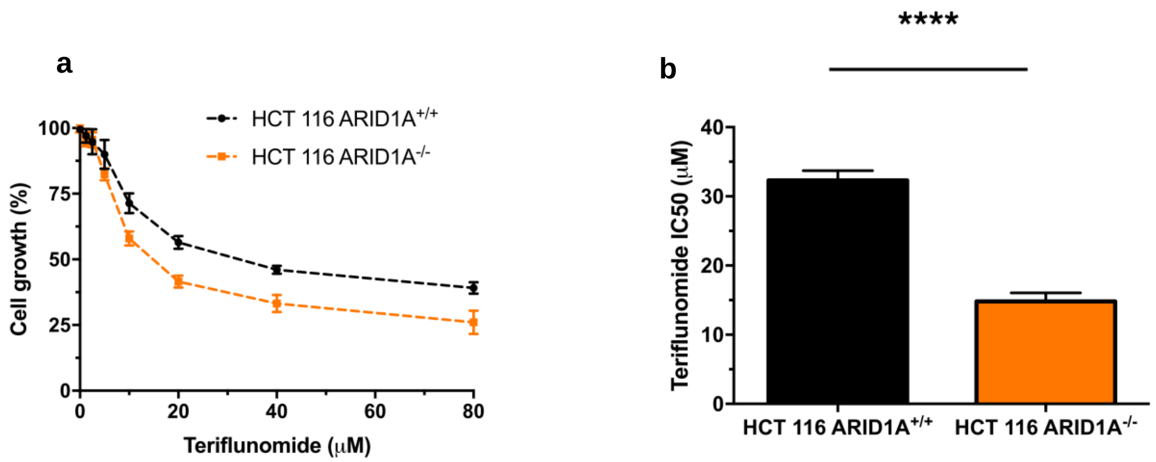


Figure S5. HCT116 *ARID1A* knockout cells are more sensitive to teriflunomide compared to control cells (HCT116 *ARID1A* wild-type). Panel (**a**) shows a cell growth curve following 72 h of treatment with the indicated concentrations of teriflunomide. Each symbol represents the mean ± SD of 3 independent experiments. Panel (**b**) shows the teriflunomide IC50, the drug concentration corresponding to a 50% decrease in cell viability, for HCT116 *ARID1A* knockout cells compared with control cells. Each bar shows the mean ± SD of 3 independent experiments. ****$P < 0.0001$, two-tailed *t*-test.

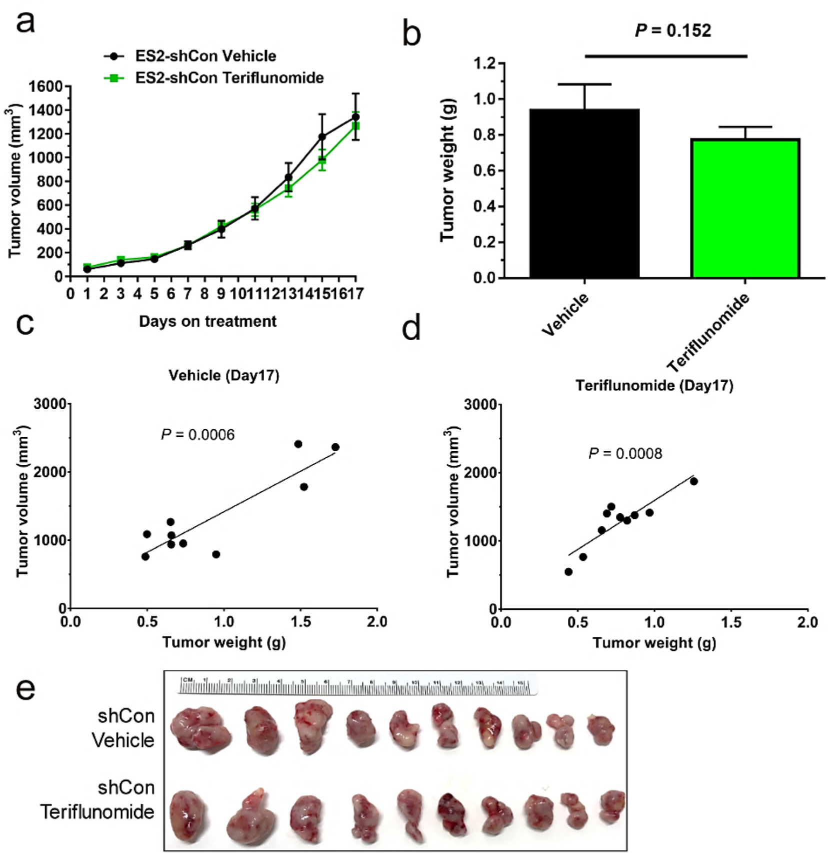


Figure S6. The effect of teriflunomide was evaluated in an ES2-shCon tumor xenograft model. **a**, The xenograft model was generated by subcutaneously injecting ES2-shCon cells in Matrigel (1:1) into athymic nude mice. Teriflunomide (4 mg/kg) or vehicle was intraperitoneally injected every other day. Tumor size was recorded on the same day. The effect of teriflunomide on tumor xenograft growth is shown by depiction of the mean tumor volume ± s.e.m. (N = 10 animals/group) for the following two groups: shCon treated with vehicle (black line) and shCon treated with teriflunomide (green line). Teriflunomide showed no effect between the two groups. **b**, the terminal tumor volumes on day 17 is summarized by graphing the tumor weight. **c** and **d**, Analyses of the correlation of tumor weight with tumor volume are shown. **e**, Terminal tumor images on Day 17 are shown.

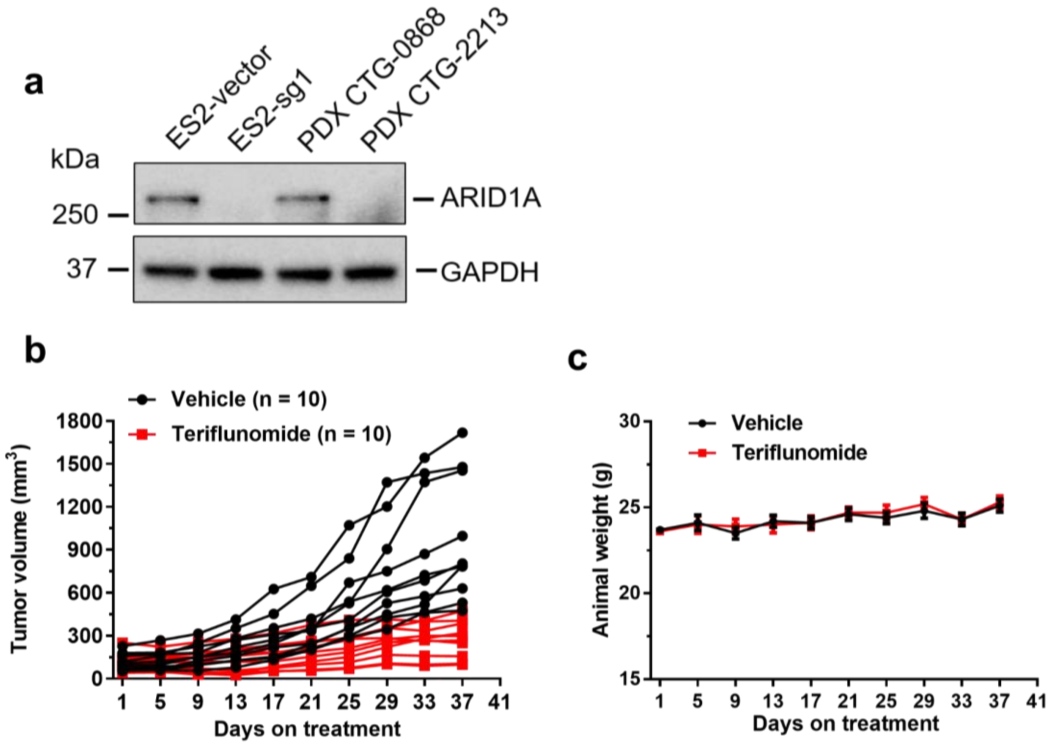


Figure S7. The effect of teriflunomide in PDXs model. **a**, ARID1A protein expression in PDX tumor samples were analyzed by western blot assay. *ARID1A* wild-type ES2-vector and *ARID1A* knock-out ES2-sg1 groups showed as positive and negative controls, respectively. PDX CTG-2213 tumor, which was ARID1A deficient tumor, was used in our *ARID1A* deficient PDX model. **b**, Individual tumor growth in both the treatment and vehicle control groups. **c**, There was no significant weight loss or toxicity in mice in either the treatment or vehicle control group. Means ± s.e.m. were calculated in each group.

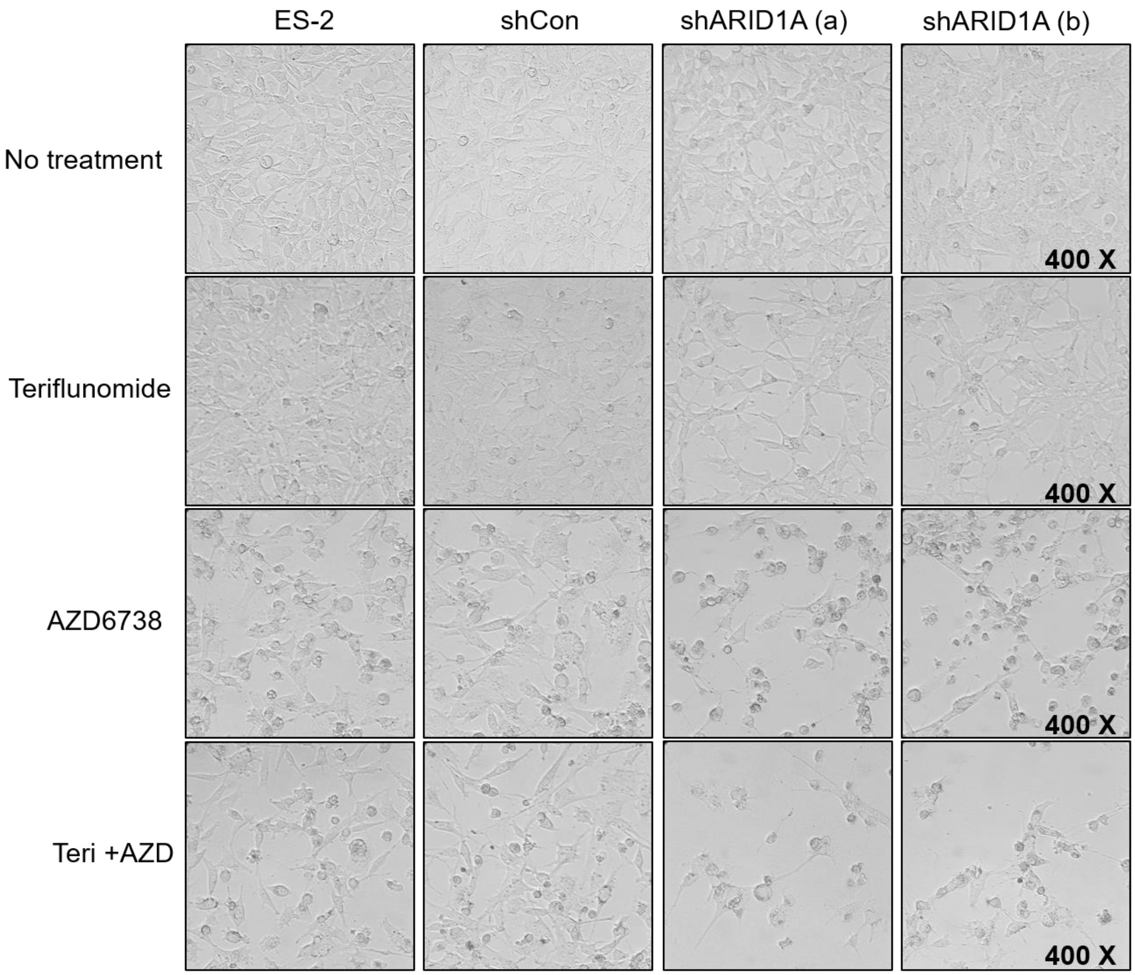


Figure S8. Cell growth arrest caused by combination treatment of ES2 cells with teriflunomide and AZD6738 for 72 h is quantified. The doses of 15 µM teriflunomide and 1.5 µM AZD6738 were chosen based on their $IC_{50}$ in ES2 cells. *ARID1A*-knockdown cells (shARID1A) showed cell growth arrest in the single-drug treatment groups, but the greatest effect was in the Teri+AZD combination treatment group.

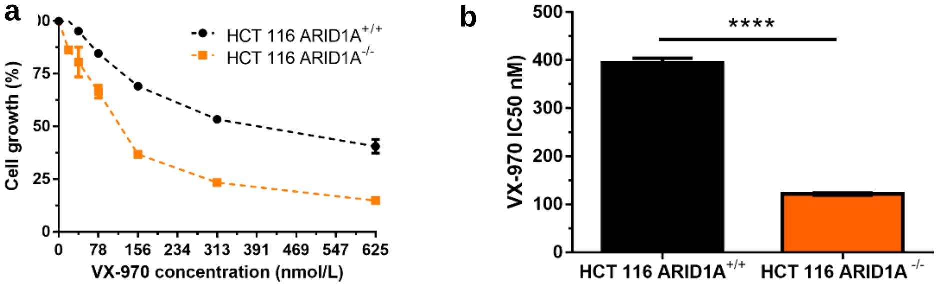


Figure S9. *ARID1A*-knockout HCT116 cells were more sensitive to VX-970 compared to control cells. **a**, cell growth curve following 72 h of treatment with the indicated concentrations of VX-970 is shown. Each symbol represents the mean ± SD of three independent experiments. **b**, the $IC_{50}$, or the drug concentration corresponding to a 50% decrease in cell viability, for VX-970 treatment of *ARID1A*-knockout HCT116 cells compared with control cells is depicted. Each bar shows the mean ± SD of three independent experiments. ****$P < 0.0001$, two-tailed t-test.

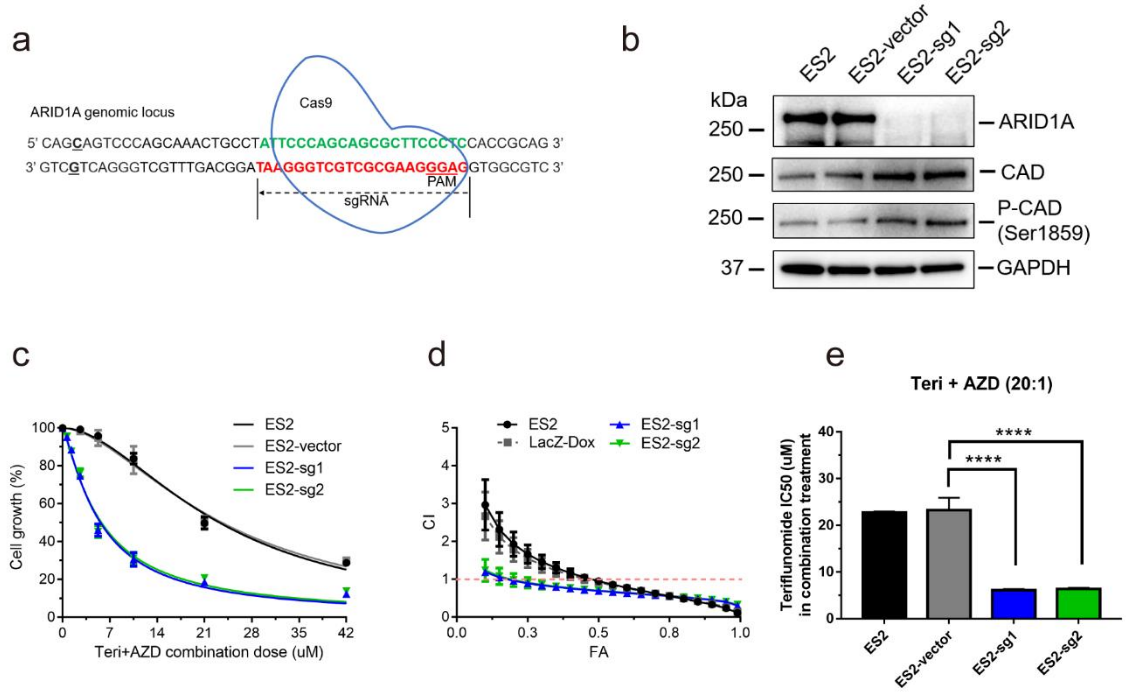


Figure S10. *ARID1A* knockout ES2 cells were more sensitive to combination treatment of teriflunomide with AZD6738. **a**, schematic of *ARID1A* knockout in ES2 cells using CRISPR-cas9 system. **b**, The absence of ARID1A protein expression by Western blot. Increase of CAD and P-CAD Ser1859 in *ARID1A* knockout cell clones. **c**, the effect of combination treatment of ES2 cells with teriflunomide and AZD6738 (Teri : AZD = 20:1) for 72 h was quantified. ARID1A-knockout cells, depicted by the blue and green lines, were even more sensitive to combination treatment compared to non-ARID1A-induced cells. **d**, the data shown in (**c**) are summarized by graphing FA-CI plots, with x = fraction affected (FA) vs. y = combination index (CI) (the Chou-Talalay plot). The FA-CI curves showed the detail synergic effect at each drug concentration. CI < 1, = 1, and > 1 indicate synergism, an additive effect, and antagonism, respectively. Detailed analyses of the combination treatments in (**c**), is shown in Supplementary Tables S8. **e**, the data shown in (**c**) was summarized by graphing the teriflunomide concentration of combination treatment that results in a 50% growth inhibitory effect (IC50). The bars depict the mean IC50 ± SD for five independent experiments. Differences in IC50 were evaluated using one-way ANOVA with Tukey's post-test; ****$P$ < 0.001.

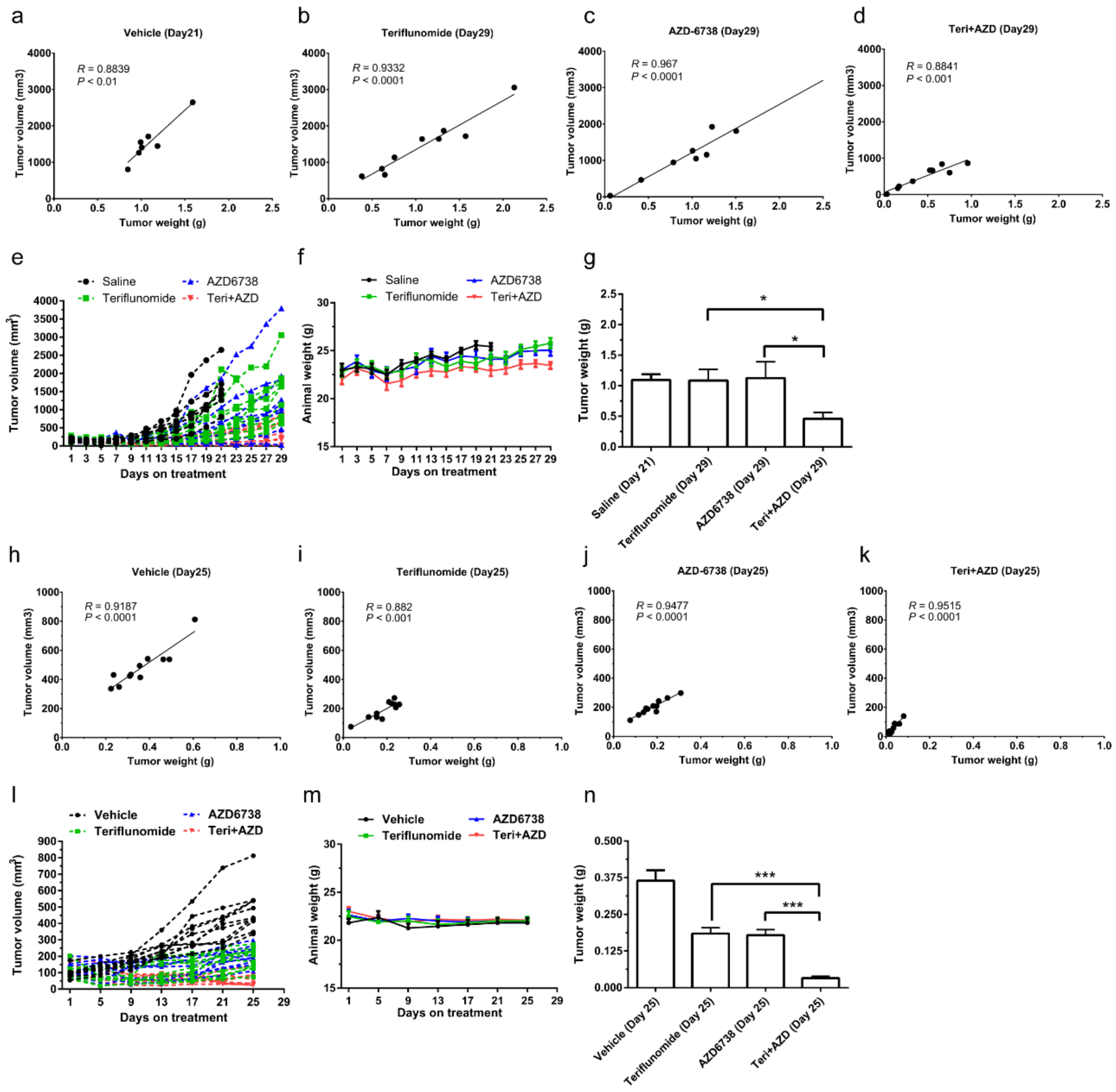


Figure S11. Combination treatment with DHODHi and ATRi *in vivo*. ES2-shARID1A xenografts (**a-g**) and PDXs (**h-n**) were treated with vehicle control, teriflunomide, AZD6738, or teriflunomide plus AZD6738. **a**-**d** and **h**-**k**, correlation analysis of tumor weight with tumor volume. High correlation of tumor weight with tumor volume in each xenograft treatment group. **e** and **l**, Individual tumor growth in all four groups. **f** and **m**, animal body weight growth in all four groups. **g** and **n**, average tumor weight comparison in ES2-shARID1A xenografts and PDXs. *$P < 0.015$, ***$P < 0.001$. Means ± s.e.m. was calculated in each group.
